# Mapping human pre-rRNA processing and modification at single nucleotide resolution using long read Nanopore sequencing

**DOI:** 10.1101/2025.03.01.640970

**Authors:** Stefan Pastore, Ludivine Wacheul, Lioba Lehmann, Stefan Mündnich, Beat Lutz, Mark Helm, Susanne Gerber, Denis L.J. Lafontaine, Tamer Butto

**Affiliations:** Institute of Pharmaceutical and Biomedical Sciences, Johannes Gutenberg-University Mainz, Mainz 55128, Germany; RNA Molecular Biology, Fonds de la Recherche Scientifique (F.R.S./FNRS), Université libre de Bruxelles (ULB), Biopark campus, B-6041 Gosselies, Belgium; Institute of Human Genetics, University Medical Center of the Johannes Gutenberg University Mainz, Mainz 55131, Germany; Leibniz Institute for Resilience Research (LIR), 55122 Mainz, Germany; Institute of Physiological Chemistry, University Medical Center Mainz, 55128 Mainz, Germany

**Keywords:** NanoRibolyzer, ribosome biogenesis, nucleolus, pre-rRNA intermediates, pre-rRNA processing, fingerprinting, Nanopore-sequencing, pseudouridines

## Abstract

Ribosome biogenesis requires the synthesis and processing of precursor rRNAs (pre-rRNAs) into mature rRNAs. Traditional methods like northern blotting and metabolic labeling offer limited resolution. We present NanoRibolyzer, a nanopore-based, long-read sequencing approach that enables ab initio identification and quantification of rRNA precursors. Using both supervised and unsupervised mapping, it detects known and novel pre-rRNA species and defines cleavage sites at single-nucleotide resolution. A simple cell fractionation protocol provides spatial separation of nuclear and cytoplasmic pre-rRNAs. Targeted knockdowns quantify intermediate accumulations, revealing condition-specific processing ‘fingerprints’ with biomarker potential. Pseudouridine mapping shows that the primary 47S transcript is extensively modified, while aberrant products are not. With its high resolution and unique mapping strategy, NanoRibolyzer offers new insights into rRNA processing and modification, enhancing our understanding of ribosome biogenesis.

## Introduction

Ribosomes are ribonucleoprotein nanomachines responsible for protein synthesis in all living cells^1,2^. Ribosome biogenesis is a complex process involving the synthesis, processing, and modification of precursor ribosomal RNAs (pre-rRNAs), as well as RNA folding and packaging into functional ribosomal subunits. In eukaryotes, this pathway is initiated in the nucleolus, where a large ribosomal RNA precursor (pre-rRNA), the 47S, is synthesized by RNA polymerase I (Pol I)^3,4^. The 47S contains sequences for three out of four rRNAs (the 18S, 5.8S, and 28S) interspersed with non-coding spacers (**Fig 1A and S1**). The fourth rRNA, 5S, is produced independently by Pol III, in the nucleoplasm. Following transcription, pre-rRNAs undergo a series of maturation steps, including processing (cleavage), modification, and packaging with ribosomal proteins, to release the mature rRNAs and produce the ribosomal subunits, which are ultimately exported to the cytoplasm where they engage in translation^5^. Throughout this multistep process, the nascent transcripts undergo extensive processing by endonucleases performing precise cleavages within the external and internal transcribed spacers (ETS and ITS, respectively) often followed by exonucleases that progressively trim pre-rRNAs, ultimately releasing the mature rRNAs (**Fig 1A and S1**). The progressive trimming of pre-rRNAs by exonucleases results in the production of transient, metastable species that likely remain largely uncharacterized. Additionally, these processes contribute to the generation of poorly defined RNA ends, further highlighting the complexity and incomplete understanding of pre-rRNA processing. Disruptions occurring at any stage of the pathway may activate regulatory cascades, including surveillance leading to the accumulation of distinctive intermediates, which can significantly impact ribosome function, cellular protein synthesis, and overall cellular homeostasis^4,6^.

**Fig 1:**
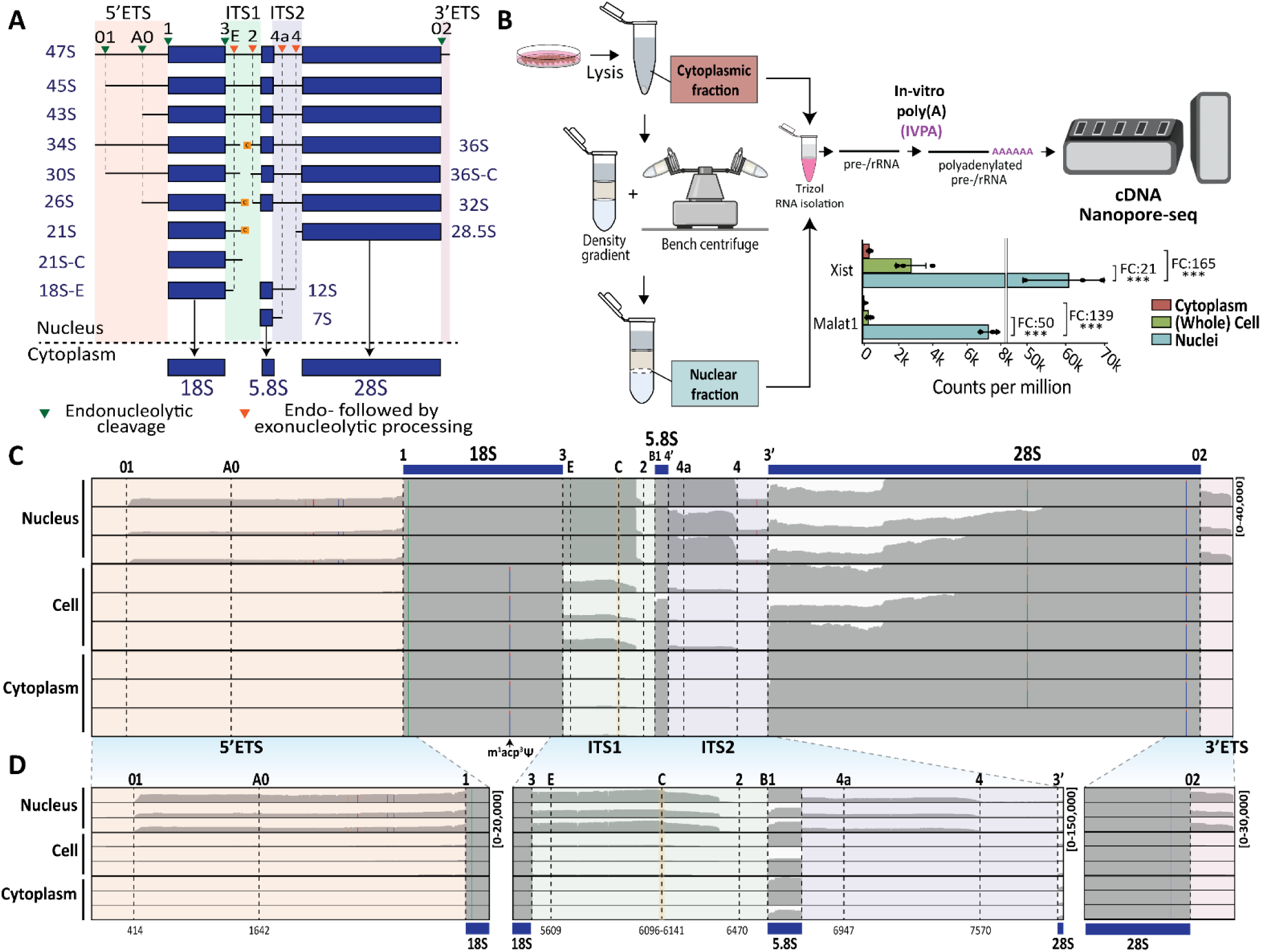
Streamlined nuclei isolation procedure utilized for isolating pre-rRNA. **A,** Simplified pre-rRNA processing pathway in human cells. The 5’ external transcribed spacer (5’ ETS) contains two primary processing sites (01 and A0). Internal transcribed spacer 1 (ITS1) has three main processing sites (3, C, and 2), while internal transcribed spacer 2 (ITS2) has at least two sites (4a and 4). The 3’ external transcribed spacer (3’ ETS) is also shown. The boundaries for the mature rRNA components are indicated as follows: 18S rRNA (sites 1 and 3), 5.8S rRNA (sites B1 and 4’), and 28S rRNA (sites 3’ and 02). The precursors for 18S rRNA biogenesis include, 30S, 26S, 21S, 21S-C, and 18S-E, while the precursors for 28S and 5.8S rRNA include, 32S, 28.5S, 12S, and 7S (Ref.^8,9,10^). The 36S, 36S-C, and 34S RNAs are associated with ribosome biogenesis perturbations. **B,** Nuclei isolation procedure for isolation of nuclear and cytoplasmic RNA followed by cDNA Nanopore-seq. (See Material and methods and supplementary Fig 1 for details). Normalized read count per million of *Xist and Malat1* transcripts in cytoplasmic, whole cell, and nuclear fractions (n=3). Fold change (FC) between the conditions is shown above each comparison. One-way ANOVA followed by Tukey test for multiple comparison post hoc test, ***p < 0.001. **C,** IGV coverage profiles of representative nuclear, whole cell and cytoplasmic samples across 47S. Data range was normalized to 40,000 across all samples to visualize the coverage profiles within the selected regions. The m¹acp³Ψ site is shown below as a representative example of a typical systematic error observed in IGV profiles. Positions with a mismatch frequency higher than 0.2 are colored. **D,** Zoom in IGV coverage profiles across 5’ ETS and 18S (left), ITS1, 5.8S and ITS2 (middle) and 28S and 3’ ETS (right) of representative nuclear, whole cell and cytoplasmic samples across 45SN1. Data range is shown on the right of each figure and was normalized across all samples to visualize the coverage profiles within the selected regions.

More than two decades of research have seen significant progress in identifying major discrete processing sites and pre-rRNA intermediates, which serve as critical markers for studying the efficacy of ribosome biogenesis (**Fig 1A and S1**). Knowledge of these intermediates has been particularly valuable for investigating aberrant precursor production that arises during processing perturbations^2^. Conventional approaches for analyzing rRNA processing intermediates, such as northern blotting, metabolic labelling, or primer extension, allow for the identification of these accumulated precursors and cleavage sites, respectively^6,8,9,10^. However, these assays require important input material (often µg range), have limited resolution and throughput. Additionally, studying pre-rRNA processing by sequencing has remained challenging due to the highly repetitive nature of rDNA arrays and poor genome annotations making it difficult to accurately map reads and distinguish between individual rDNA copies, especially with short-read sequencing technologies^11,12,13,14^.

Nanopore sequencing (nanopore-seq) has emerged as a promising technology to investigate ribosome biogenesis^15,16^. The key advantage of nanopore-seq lies in its ability to sequence long reads, such as cDNAs, as well as native RNA molecules via direct RNA sequencing (DRS), allowing for the investigation of entire transcripts including their modifications ^17,18^. These capabilities are particularly valuable for studying rRNA intermediates, which can vary significantly in length and abundance, and possibly modification levels. To this date, there are no tools that exploit long-read sequencing for the analysis of human ribosomal RNA precursors.

Here, we present NanoRibolyzer, a method that integrates state-of-the-art long-read nanopore sequencing with advanced bioinformatics to achieve spatially resolved, single-nucleotide analysis of pre-rRNA intermediates. Using a streamlined nuclei isolation protocol, we systematically profile precursor and mature rRNA species in both nuclear and cytoplasmic compartments. We further introduce precursor-specific modification analysis, uncovering the spatio-temporal dynamics of rRNA modifications. Overall, NanoRibolyzer enables comprehensive detection and quantification of both known and novel processing intermediates, including cleavage events generated by endo- and exoribonucleolytic activity, as well as pseudouridine modifications.

## Results

### A simplified nuclei isolation procedure to characterize pre-rRNA precursors

Ribosome biogenesis initiates in the nucleolus, a multiphase biomolecular condensate that resides within the nucleus. To define spatially pre-rRNA processing, we first focused on isolating highly purified nuclei fractions, from which we extracted RNA for sequencing. We developed a straightforward isolation protocol involving density gradient separation using a simple benchtop centrifuge (**Fig 1B and S2A-B**). We applied the protocol to HEK293 cells to produce highly purified nuclear and cytoplasmic RNA fractions, and as a control, whole cell total RNA (‘whole cell’) (**Fig 1B and S2A**).

After separation, a quality assessment of isolated nuclei was conducted using DAPI staining, revealing debris-free and round intact nuclei^19^ (**Fig S2C**). RNA was extracted and electropherograms produced using a TapeStation. As expected, the whole cell and cytoplasmic fractions displayed the abundant mature 18S and 28S rRNAs (all analysis performed in triplicate throughout this work, R1-R3) (**Fig S2D**). In contrast, the nuclear fractions exhibited higher molecular weight species, at the expected size for pre-rRNAs (**Fig S2D**).

Since Nanopore-based RNA library preparation strategies typically capture poly(A)+ RNA^20^, we applied an *in-vitro* polyadenylation strategy followed by long-read cDNA sequencing (**Fig 1B**). To assess enrichment of transcripts in the nuclear fractions, we quantified two nuclear long non-coding RNAs (lncRNAs): XIST and MALAT1 (**Fig 1B**, **Supplementary Table S1**). As expected, comparative analysis of the abundance of these transcripts across different cellular compartments revealed several fold-change enrichments in the nuclear fraction relative to the cytoplasm (up to 165-fold for XIST).

Having established an efficient nucleo-cytoplasmic purification protocol, we developed NanoRibolyzer, a method relying on long-read nanopore sequencing in combination with a bioinformatic pipeline specifically designed for the detection and quantification of rRNA intermediates (**Fig S3A**).

NanoRibolyzer aligns long-reads to a single 47S template (equivalent to 45SN1; GeneID:106631777), in contrast to the multiple templates found in genome annotations, offering a clearer representation of reads associated with pre-rRNA compared to whole-genome alignment. We generated an average of ∼3 million reads per sample with an average ∼75% alignment rate to the 47S template. For reference and to initiate precursor quantification analysis, we retrieved the positions of known processing sites and major precursors from literature^8,9,10^ (**Fig 1A and S1**).

We applied NanoRibolyzer to unperturbed HEK293 cells and characterized reads mapping to rRNA precursors and non-coding spacers (5’ ETS, ITS1, ITS2 and 3’ ETS) for the nuclear, cytoplasmic, and whole cell fractions (**Fig 1C-D**).

As expected, the coverage profiles revealed that the nuclear fraction had higher coverage across the ETSs and ITSs compared to the cytoplasmic ones, with the whole cell fraction displaying an intermediate coverage level (**Fig 1C-D**). The start and ends of these reads were closely aligned with annotated processing sites, such as sites 01,1, 2, 4, etc. (see Refs^8,9,10^).

These observations agree with the notion that the nuclear fraction predominantly contains ribosomal intermediates, reflecting that most steps of ribosome biogenesis occur in the nucleolus. In these initial analyses, we often observed that whole cell fractions were largely redundant with cytoplasmic fractions (see **Fig 1D**, zoomed IGV), therefore in the remainder of the manuscript we will only compare the nuclear and cytoplasmic fractions. We also noted that the hyper modified nucleotide 1-methyl-3-α- amino-α-carboxyl-propyl pseudouridine (m^1^acp^3^Ψ) at position 1248 on the 18S was clearly detected in whole cell and cytoplasmic fractions but not in nuclear ones (**Fig 1C**, arrow), which is compatible with the idea that this modification is completed in the cytoplasm^21^.

### A novel mapping strategy to characterize rRNA precursors

After confirming that the nuclear fraction coverage profiles encompass the spacer regions, we employed two complementary strategies to quantify pre-rRNA and identify processing sites: a classical supervised (template-based) and an original unsupervised (template-free) approach (**Fig S3A-C)**.

The supervised (template-based) approach considers known discrete pre-rRNA intermediates as well as the spacer regions, quantifying their relative abundance through reciprocal overlap maximization (**Fig 2A and S3B,** for details). These intermediates were derived from the literature using the positions of processing sites on the 47S, allowing us to generate the major precursors depicted in **Fig 1A** (Refs^1,2^). The quantified data can be displayed using an intuitive heatmap, showing averaged log10 reads per million (**Fig 2B and S4A**), or a histogram (**Fig S4B**), depicting the averaged reads per million for each condition.

**Fig 2.**
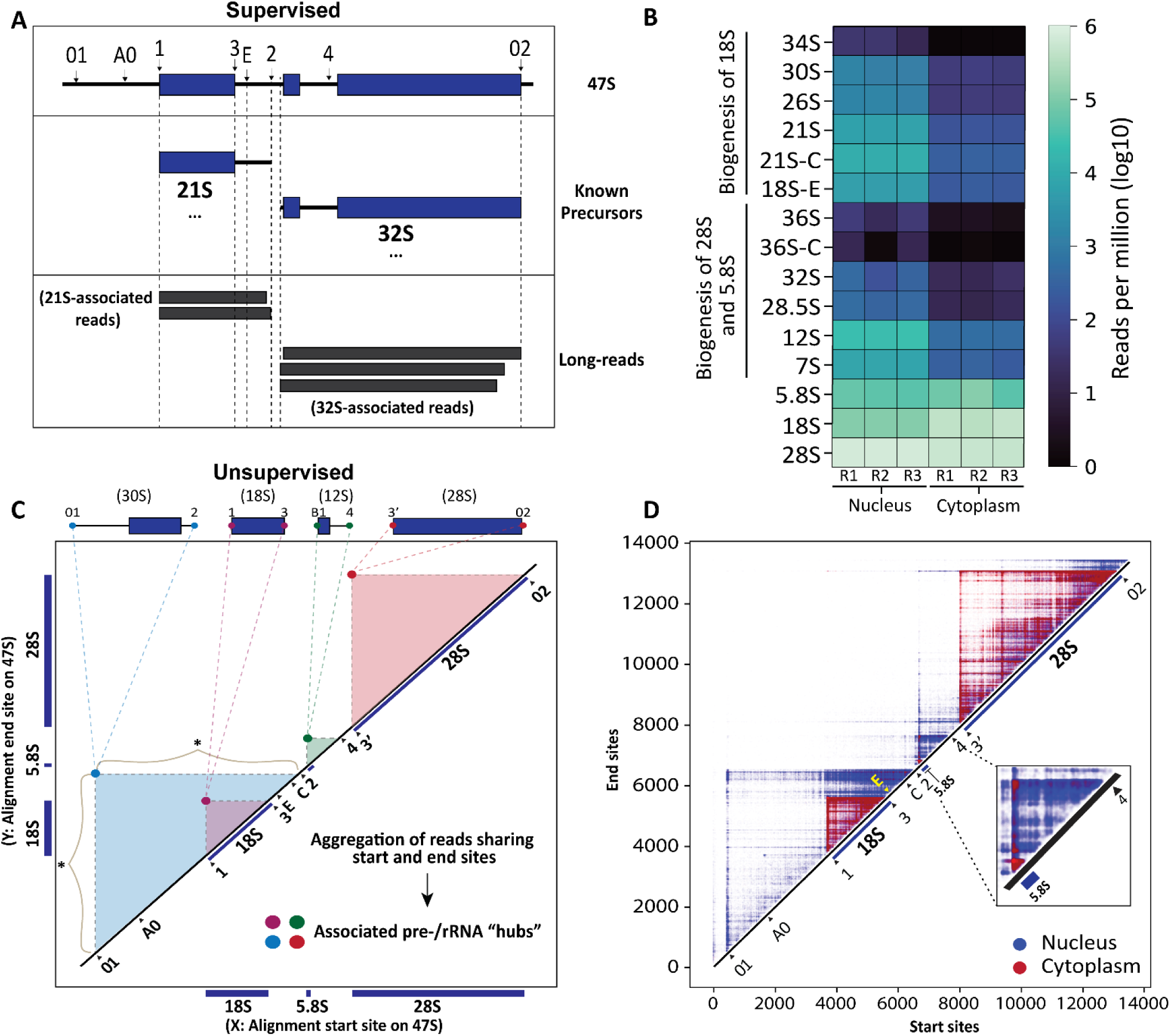
Quantification of rRNA precursors and processing sites using NanoRibolyzer. **A,** Simplified overview of supervised (template-based) approach using minimal reciprocal overlap (MRO). Query reads are compared to literature-based intermediates (Fig 1A), and each read is assigned to the intermediate with the highest overlap based on alignment start and end positions. In the example shown, the query reads are closely associated with 21S and 32S precursors. After processing all the reads, the data is presented as a relative quantification score in reads per million, allowing for clear visualization and comparison. See more details in **Supplementary Figure S3 and S4**. **B,** Quantification of detected pre-rRNA intermediates and mature rRNAs in the Nucleus and Cytoplasm (n=3). **C,** Simplified illustration of the unsupervised (template-free) approach. A 2-D matrix representing the RNA45SN1 template is constructed, with each rRNA read plotted by its start (x-axis) and end (y-axis) positions. This approach maps transcript boundaries and highlights intensity "hubs", which indicate abundant rRNA products near mature rRNA or putative processing sites. In the example shown, the intensity hubs for 30S, 18S, 12S and 28S correspond to reads clustered at the start and end of the respective pre-/rRNAs. The * symbol denotes ’processing smears’ where a stable start or end site is accompanied by multiple exonucleolytic events at the opposite end of the rRNA product. See more details in **Figure S3C. D,** Overlayed intensity matrices of nucleus (blue) and cytoplasm (red), highlighting contrasting read distributions: ETS and ITS-associated reads dominate in the nucleus (in blue), while the cytoplasm predominantly contains mature rRNA reads (in red).

Using the supervised approach, as expected, we observed that most precursors are more abundant in the nuclear fraction than in the cytoplasmic one (**Fig 2B and Fig S4A-B,** for statistical analysis, **Source Data 1**). This also illustrates that some precursors, which were historically associated with ribosome biogenesis perturbations, are indeed not produced or only marginally in unperturbed cells (e.g. 34S, 36S, 36S-C). Thus, NanoRibolyzer offers an efficient and quantitative approach to determining the relative levels of all major pre-rRNA precursors and mature rRNAs.

Having mapped long read sequences to specific pre-rRNA precursors and mature rRNAs, our next goal was to identify precisely their prevalent start and end sites, defining processing sites. To this aim, we computed significant starting and ending sites using BAM files from the template-based data and determined their abundance using the sequence 45SN1 (hg38), as a reference. Starting and ending sites occurring at least two standard deviations above the mean were considered significant, and results were stored in TSV and BED file formats for visualization (**Fig S5A-D**). Importantly, we only kept sites which were shared in at least two samples of our triplicates to identify the most significant ones and quantify their prevalence across the 47S sequence (**Source Data 1**).

An initial inspection of the cleavage sites revealed consistent positions associated within spacer regions and the start and end points of mature rRNAs **(Fig S5B**, start sites in green; end sites in orange). Cleavages detected within spacer regions, particularly those located near previously described processing sites were considered as *bona fide* processing sites (**Fig S5B-D, see Fig 1A and S1**, in specific cases, their resolution had not been resolved to a single nucleotide). In contrast, cleavages within mature rRNA sequences may correspond to aberrant cleavages producing by- products destined for degradation **(Fig S5B).** As expected, in the cytoplasm, where pre-ribosomes are nearly completed or mature ribosomes engaged in translation or stored, cleavage sites were restricted to the ends of mature rRNA (**Fig S5B).** In stark contrast, nuclear samples exhibited prevalent cleavages within the spacer regions reflecting ongoing maturation **(Fig S5B-D).** Overall, our supervised approach can provide both quantitative measurement of the precursors as well as information about the prevalent cleavage sites which correspond to processing sites.

Next, we developed an original unsupervised (template-free) approach providing an unbiased mapping method. The rationale for implementing it was to study processing steps that involve progressive exonuclease trimming of pre-rRNAs generating transient, metastable species that are often poorly characterized as they produce ill-defined ends (**Fig 1A** and **S1**). In this case, reads are plotted in a 2-D graph, forming a matrix (**Fig 2C and S3C**). The x-axis and y-axis span from the transcription start site to the termination site of the primary transcript. Each rRNA read is plotted onto the matrix based on its starting and ending positions (x and y coordinates, respectively), enabling precise mapping and analysis of transcript boundaries (**Fig 2C and S3C**). Such displays offer an intuitive visualization of intensity "hubs" representing abundant rRNA products whose ends are at, or close to, well-established or putative novel processing sites (**Fig 2C and S3C**, see methods for further detail).

We applied the unsupervised approach to the nuclear and cytoplasmic fractions of unperturbed HEK293 cells to identify the unbiased aggregation of reads associated with the 47S template. In the nuclear fraction, we observed an accumulation of reads associated with the ETS and ITS regions, correlating with processing sites and depicted as blue “pyramids” (**Fig 2D**). For example, striking pyramids between site 01 in the 5’ ETS and site 2 in ITS1, defining precursors of 18S (see below), or between the 5’ end of 5.8S and site 4 in ITS2, representing 3’ extended precursors of 5.8S destined to be processed 3’-5’ by the multi-subunit RNA exosome^6,22^ (**Fig 2D**, zoomed in inset, **and S4C**). This approach also enabled the detection of previously unobserved low-abundance and metastable processing intermediates, as well as by-products, appearing as “processing smears” in the intensity matrix (**Fig 2C**, see asterisks and **Fig S6A-B**).

In contrast, reads in the cytoplasmic fraction (in red in **Fig 2D**) were nearly exclusively associated with the three mature rRNAs, the 18S, 5.8S and 28S (**Fig 2D**, see inset for 5.8S). The individual matrices for the three replicates are shown in **Fig S4C**, attesting to the robustness of the analysis.

In conclusion, by combining supervised and unsupervised mapping approaches, we not only leverage existing knowledge of abundant and long-lived pre-rRNA precursors, previously mapped using conventional techniques, but also create new opportunities for identifying previously undetected, low-abundance, metastable processing intermediates and by-products.

### Redefining pre-rRNA processing sites to single nucleotide resolution

Next, we followed an agnostic approach to remap the processing sites by analyzing the intensity matrix obtained by unsupervised mapping. To achieve this, we extracted approximate coordinates from detectable intensity “hubs” in the nuclear fraction and annotated the associated 5’ and 3’ processing sites (**Fig 3A**). These coordinates are considered estimates due to some variability among minor RNA species end points within a hub, thus we bioinformatically selected the most predominant ones in proximity to putative processing sites using a threshold of >10 reads per hub.

**Figure 3.**
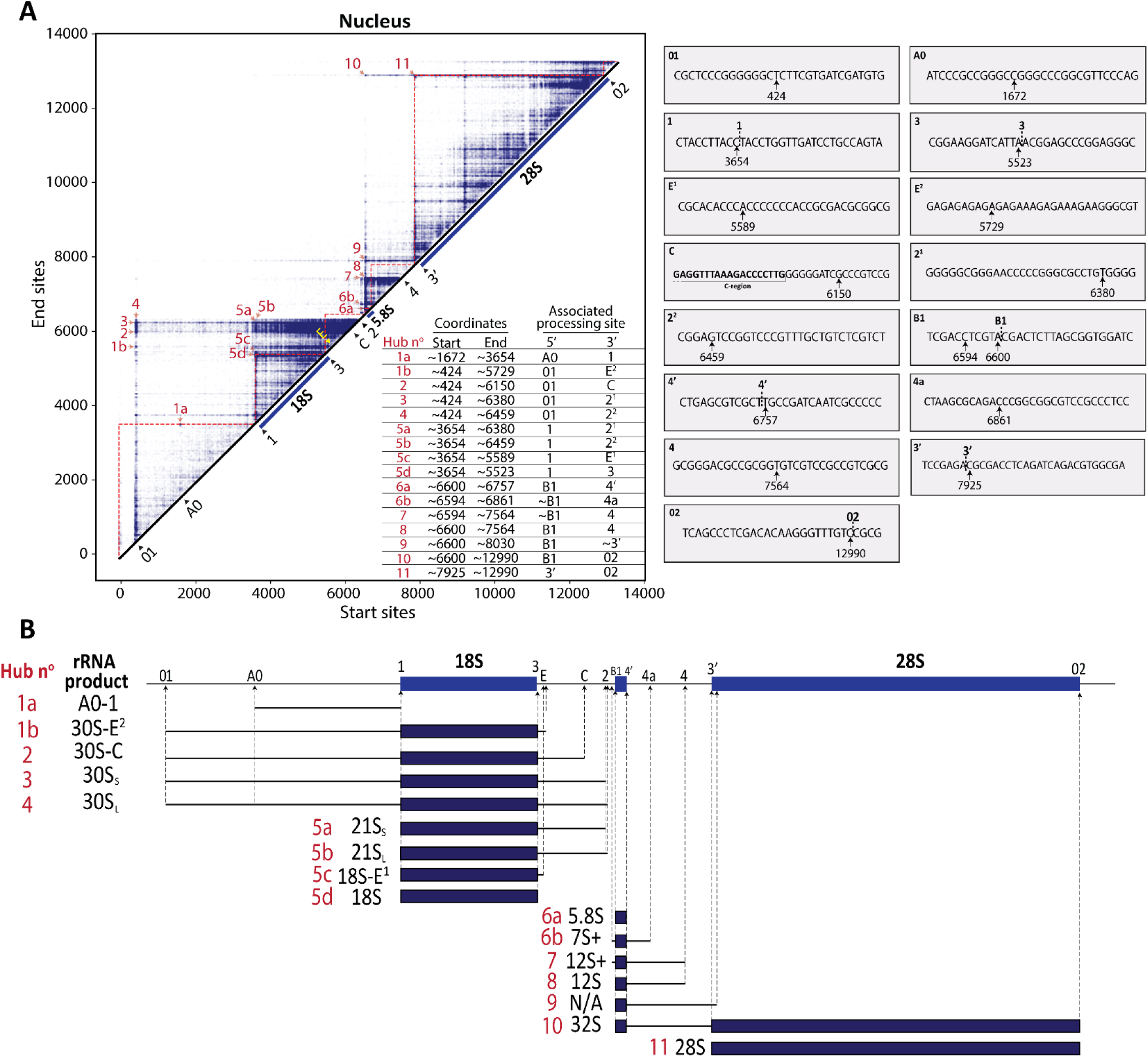
Identification of processing sites and novel precursors with nucleotide resolution. **A**, Left, Intensity matrix of nuclear conditions, with highlighted intensity “hubs” numbered in red. The “hubs” were identified based on intensity and closeness to *bona fide* processing sites. Right, The insets provide the of each mapped processing sites. Note that “hubs” exhibiting vertical conservation at the 5’ end indicates 3’-5’ exonuclease activity, while horizontal conservation at the 3’ end suggests 5’-3’ exonuclease activity (**See Figure S5A-B**). **B,** Illustration of the identified rRNA products in nucleus, as derived from the intensity hubs table shown in 3A. The corresponding rRNA products are shown on the left, alongside the site number (red) and associated precursor (black). In hubs number 6b and 7, the + symbolizes the 5’ extension upstream 5.8S sequence.

Analysis of the intensity matrix revealed obvious processing sites associated with several known cleavages, allowing to map them to single nucleotide resolution and to show, for some of them, they may in fact correspond to two cleavages. We will now review them briefly, one by one:

1. For **site 01**, the start site of the associated intensity hub maps to position C^424^^T (where the proposed cleavage site is marked by ^), which confirms previous findings^23^ (**Fig 3A**; **Hubs n° 1b, 2, 3, and 4**).
2. For **site A0**, we identified a hub with a start site at ^1672^C^G, located 20 nucleotides downstream of the proposed processing site^24^ (**Fig 3A**; **Hub n° 1a)**.
3. For **site 1**, marking the 5’ end of 18S, we detected multiple hubs containing the start site ^3654^C^T (**Fig 3A**; **Hubs n° 5a, 5b, 5c, and 5d)**
4. For **site 3**, corresponding to the 3’ end of 18S, we confirmed site ^5523^A^A (**Fig 3A**; **Hub n° 5d).**
5. For **site E**, interestingly, we identified two distinct end points: one we refer to as “**E^1^**”, located 11 nucleotides downstream of the previously reported site E (Refs ^25,26^), with multiple cleavages observed at coordinate C^A^5589^, and the other we refer to as “**E^2^**”, located in an AG-rich region, with reproducible end cleavages around G^A^5729^ (**Fig 3A and S6A-B ; Hubs n° 1b and 5c)**.
6. For **site C**, we identified an end point located at site C^G^6150^, 8 nucleotides downstream of the previously described conserved “region C” (Ref.^25^) (**Fig 3A**; **Hub n° 2**).
7. For **site 2**, our analysis also revealed two distinct intensity hubs: site “**2^1^**”, located at G^T^6380^, and site “**2^2^**”, positioned near A^G^6459^, in agreement with the former observations ^27,28^ (**Fig 3A and S6A-B; Hubs n° 3, 4, 5a, and 5b**). These sites could also be observed in the coverage profiles upstream the putative site 2 (**Fig 1C-D**). These findings reveal two potential precursors associated with 21S, as previously suggested^29^.
8. For site **B1**, we detected the intensity hub precisely at the expected 5’ end of 5.8S, at ^5600^A^CGA, defining the short form of 5.8S, the 5.8SS (**Fig 3A**; **Hub n° 6a, 8, 9, and 10)**. Interestingly, we identified an intensity hub six nucleotides upstream of the 5.8S start site at position ^6594^C^C (**Fig 3A**; **annotated as ∼B1 in hubs 6b and 7**). This site most likely corresponds to the previously described 5’ extended version of 5.8S (both coexist in cells, see e.g. ^30^), defining the long form of 5.8S, the 5.8SL (^8,30,31^).
9. For **site 4’**, corresponding to the end site of 5.8S, we confirmed the end point located at site _T^T_6757.
10. For the still poorly characterized **site 4a** in ITS2 (Refs ^8,9,10^), we reveal a very clear end point intensity hub located at AGA^C^6861^ (**Fig 3A**; **Hub n° 6b**).
11. For the uncharacterized **site 4**, we identified a distinct intensity hub at the end coordinate T^7564^GT, defining its cleavage site (**Fig 3A**; **Hubs n° 7-8)**.
12. Next, at **site 3’**, marking the start of 28S, we confirmed the start point located at site ^7925^A^C (**Fig 3A**; **Hub n° 11)**.
13. And lastly, for **site 02**, marking the end of 28S, we confirmed the end point located at site^12990^C^C (**Fig 3A**; **Hub n° 11)**.

Many of the identified cleavage sites align with processing sites detected via the supervised approach, further validating accuracy **(Fig S5B-D)**, and globally agree with the literature adding precision.

Overall, NanoRibolyzer’s unsupervised mapping approach enables the detection and characterization of both known and novel processing sites. We mapped thirteen known processing sites at single-nucleotide resolution, revealing multiple endpoints at two sites (E^1^ and E^2^, and 2^1^ and 2^2^).

Having shown the intensity hub mapping strategy identifies novel processing sites, we next considered the novel precursors they define (**Fig 3B**). Typically, several intensity hubs correspond to species starting at site 01 and ending at different sites, such as site 3 (18S 3’ end), E, C, 2^1^ and 2^2^ (**Fig 3B**; **Hubs n° 1b and 2-4**). The detection of such precursors illustrates that cleavage in ITS1 can occur prior to cleavage at site A0, at least to some extent, in unperturbed cells. We also identified species ending at a known site but starting at unknown continuous positions (“smear”), indicating they are subject exonucleolytic digestion (**Fig S6B**). Additionally, we refined further the 21S annotation, categorizing the precursor into two distinct forms: 21S_S_ (small) and 21S_L_ (large), as previously observed^29^ (**Fig 3B**; **Hubs n° 5a-b, and S6B**). In the case of 5.8S rRNA maturation, we identified abundant 3′-extended precursors that reflect the progressive exoribonucleolytic trimming of 12S to 7S and ultimately to 5.8S. This process involves substrate handover between RNA exosome subunits^6^ (**Fig. 3B**; **Hubs no. 6b–7 and S6B**).

In conclusion, we highlight how rRNA products, including known and novel metastable precursors, can be detected and analyzed at high resolution and sensitivity with NanoRibolyzer.

### NanoRibolyzer captures pre-rRNA intermediate changes following processing perturbations

Having shown the ability to capture pre-rRNA processing in unperturbed cells, we next aimed to evaluate how efficiently the method can detect processing perturbations. To achieve this, we selected key processing factors involved in the maturation of each spacer sequence and depleted them (**Fig 4A**).

**Figure 4.**
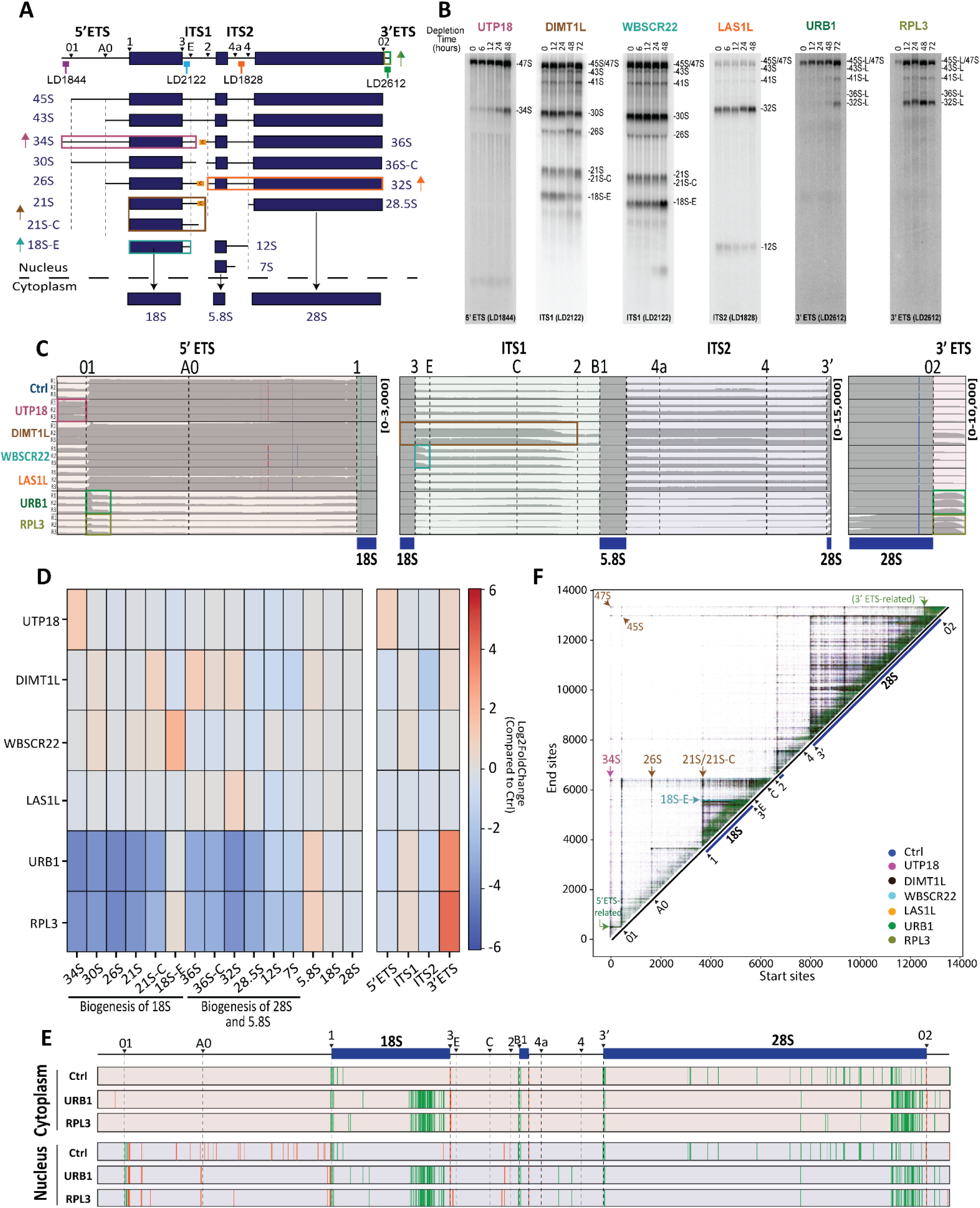
Quantification of pre-rRNAs following processing perturbations. **A**, Major effects of knockdown of key factors involved in maturation of spacer regions. Knockdowns of UTP18 (purple, within 5’ ETS), DIMT1L (brown, within ITS1 and 5’ ETS), WBSCR22 (turquoise, within ITS1), LAS1L (orange, within ITS2), URB1 (green, within 3’ ETS) and RPL3 (dark yellow, within 3’ ETS) causes the accumulation of 34S, 18S-E, 21S-C, 32S and 3’ ETS ribosomal intermediates/spacer region, respectively. Location of probes used in panel B are indicated. **B**, Assessment of intermediate accumulation following processing perturbation using northern blotting. Each target was depleted in HEK293 cells in a time course (from 6 to up to 72 h) to identify the best time point for NanoRibolyzer analysis. Based on this analysis, the following depletion time points were selected: UTP18 (48 h), DIMT1L (72 h), WBSCR22 (48 h), LAS1L (48 h), URB1 (72 h) and RPL3 (72 h). **C**, IGV coverage profiles of nuclear reads of control and knockdown of UTP18, DIMT1L, WBSCR22, LAS1L, URB1 and RPL3 across the 5’ ETS (left), ITS1, 5.8S, and ITS2 regions (middle) and 3’ ETS (left). The regions affected by the processing perturbation are highlighted in the corresponding colors shown in panel A. **D**, Log2FoldChange of quantified pre-rRNA intermediates and mature rRNAs in the nucleus of UTP18, DIMT1L, WBSCR22, LAS1L, URB1 and RPL3 knockdown samples compared to control (n=3 each). **E**, Visual representation of prevalent start (green) and end (orange) cleavage sites across the 47S rRNA in cytoplasmic (top) and nuclear (bottom) samples, comparing control samples to knockdown samples of URB1, and RPL3. Each cleavage site displayed was detected in at least two out of three replicates. **F,** Overlayed Intensity matrix of nucleus of control, UTP18, DIMT1L, WBSCR22, LAS1L, URB1 and RPL3 knockdown samples. Selected intensity hubs are indicated with arrows, displaying their coordinates (start, end) along with the associated precursor, where applicable.

For the 5’ ETS, we targeted UTP18, a component of the SSU-processome, whose depletion results in the accumulation of the aberrant 34S species (Ref.^32^). For ITS1, we selected DIMT1L and WBSCR22, whose depletion results respectively in accumulation of the 21S/21S-C and the 18S-E pre-rRNA^33,34^.

For ITS2, we selected LAS1L, whose absence causes accumulation of 32S pre-rRNA^35^,. Lastly, for the 3′ ETS, we chose URB1, recently shown to be important for 3’ ETS removal^36^ and RPL3, which is involved in 25S rRNA stabilization via 5’-3’ pre-RNA interaction in yeast^37^, but whose role in 3′ ETS processing in humans remains undefined.

First, we assessed the depletion of each factor using northern blots, confirming the expected precursor accumulation (**Fig 4B**). For NanoRibolyzer analysis, we chose the optimal depletion time point for each factor (48 h for UTP18, WBSCR22, and LAS1L, and 72 h for DIMT1L, URB1 and RPL3) (**Fig 4B**).

RNA from nuclear and cytoplasmic fractions was extracted in triplicate and analyzed using cDNA Nanopore-seq, as described above. Coverage profiles revealed distinct differences in processing factor knockdowns compared to controls (**Fig 4C and S7A-C**). The effects observed by northern blotting were all confirmed by nanopore-seq (**Fig 4B-C**).

The depletion of UTP18 led to a substantially increased coverage upstream of site 01, corresponding to the 34S species (**Fig 4C**, purple rectangle, **and S7B)**. Knocking down DIMT1L or WBSCR22 increased coverage in ITS1 downstream of the 3’ end of 18S, corresponding to 21S/21S-C (in case of DIMT1L, dark brown rectangle) and 18S-E accumulation (WBSCR22, turquoise rectangle), respectively (**Fig 4C and S7C**). Depletion of LAS1L revealed increased reads across ITS2, in agreement with 32S accumulation (**Fig 4C** and **S7C**). Both URB1 and RPL3 depleted samples showed increased coverage across the 3′ ETS (**Fig 4C and S7A**). For URB1, this confirms the accumulation of 3’ ETS extended species in URB1-deficient cells^36^. The observation upon RPL3 knockdown reveals it is also indispensable for this maturation. Each RNA species was quantified (supervised approach) and visualized as heatmaps (**Fig 4D and S8A**) or histograms (**Fig S8B-E**) confirming the observations on the IGV traces (**Source Data 2**).

In addition to confirming bold phenotypes formerly associated with the selected factors, it is interesting to note that the nanopore-seq approach revealed novel unexpected features. For example the striking, and highly reproducible accumulation of fragments encompassing part of the 5’ ETS starting at site 01 upon RPL3 and URB1 depletion (see green and dark yellow rectangles in **Fig 4C and S7B**), as well as presence of a remarkable fingerprint consisting of multiple consecutive cleavages within the 3’ end portion of the 18S and 28S sequences, which we interpret as increased turn over (**Fig 4E** and **S8F, Source Data 3).** Just as unexpected was to observe the effect of DIMT1L, a small subunit assembly factor, on 3’ ETS processing and 32S maturation (**Fig 4C-D**, and **Fig S7A** and **C**).

Lastly, using the unsupervised approach, we mapped the nuclear reads onto the intensity matrix (**Fig 4F and S9A-F**). Upon UTP18 depletion, a unique hub corresponding to 34S was detected (red arrow, **Fig 4F and S9A**). Upon DIMT1L depletion, hubs associated with 21S/21S-C and 26S were seen (brown arrows), indicating impaired processing between sites C and 2, in combination with defects in the 5′ ETS (**4F and S9B)**. Additionally, we observed notable accumulation of early precursors such as 47S and 45S, consistent with previous findings^34^. Depletion of WBSCR22 revealed accumulation of a hub associated with 18S-E (turquoise arrow), with shortened start coordinates, indicating some level of 5’-3’ degradation of 18S precursors (**Fig 4F and S9C**). Knockdown of LAS1L showed reduced aggregated reads at processing site 4 and accumulation at 32S (orange arrow, **Fig S9D**). URB1 and RPL3 knockdown exhibited accumulation of reads downstream of processing site 02 (3’ end of 28S) corresponding to 3’ ETS containing RNAs (green arrows, **Fig 4F and S9E-F)**. Additionally, we observed an increased accumulation upstream of the 01 site as observed in the coverage profiles and yellow arrows, **Fig 4F and S9E-F**) and across mature rRNA regions, likely reflecting by-product buildup and elevated turnover, as supported by cleavage site analysis (**Fig 4E and S8F**).

In conclusion, NanoRibolyzer identifies and quantifies pre-rRNAs and mature rRNAs, defining processing sites to single nucleotide, identifying novel ’fingerprints’, confirming effects of factor depletion and revealing novel aspects.

### NanoRibolyzer uncovers spatio-temporal dynamics of RNA modification

Ribosomal RNA is abundantly decorated by covalent modifications^38^. To investigate the spatial dynamics of these modifications, we employed direct RNA sequencing (DRS), a technique that sequences native RNA and detects modifications through distinct current signatures that differentiate modified from unmodified bases^40^.

First, we selected three base modifications known to occur during late 18S biogenesis: *N^7^*- methylguanosine m⁷G, deposited by WBSCR22 at G1639 (Ref.^34^), *N^6^,N^6^*-dimethyladenosine (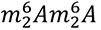) deposited by DIMT1L at A1735 and A1736 (Ref.^34^ ) and m¹acp³Ψ which is finalized by addition of the acp group by TSR3 at U1248 (Ref^39^) (**Fig S10A-C**). To obtain ground truth, we generated an *in vitro* transcribed sample (IVT 18S) to serve as constitutively unmodified negative control. Additionally, we used RNA extracted from cells inactivated for DIMT1L, WBSCR22, or TSR3 to serve as negative controls. We performed signal mapping refinement (conversion of sequence to raw signal) using *Remora* and analyzed the deviations around each modification site across late precursors (21S, 21S-C, 18S-E) and mature rRNA (18S) in both nuclear and cytoplasmic RNA.

For all three modifications, the signal corresponding to precursors closely resembled that observed in inactive mutants or IVT controls, indicating a lack of modification at these sites (**Fig S10D-F)**. In contrast, cytoplasmic 18S showed a distinct signal shift at these positions, consistent with presence of the modification on mature rRNA. For m⁷G, the shift between unmodified (IVT, inactive enzyme, and precursors) and mature cytoplasmic RNA was obvious (panel D). For 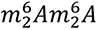, it was less obvious but visible (panel E). For m¹acp³Ψ, the situation was a bit more complex to interpret because this modification is formed in three steps: 2’-O methylation by SNORA13, methylation by EMG1, and addition of acp by TSR3. The IVT control lacking all modifications (yellow trace) and the sample extracted from cells inactivated for TSR3 lacking only the acp group (black trace) are clearly shifted from each other and from the mature fully modified cytoplasmic 18S rRNA (in red). The nuclear precursor traces (21S, 21S-C, 18S-E) superimpose nicely with the sample inactivated for TSR3, confirming earlier prediction that the addition of the acp moiety is a late event.

In conclusion, NanoRibolyzer pipeline can assign base modification status to particular pre-rRNA precursor based on raw signal analysis.

### Ribosomal RNA pseudouridine formation is mostly cotranscriptional

Next, we aimed to detect the abundant pseudouridine (Ψ), a uridine isomer installed by DKC1 guided by box H/ACA snoRNAs^40^. For this, we used first our nuclear and cytoplasmic RNA fractions and employed SeqTagger^41^ to multiplex up to four DRS barcodes per flow cell (two replicates per condition). Following sequencing, pseudouridine levels were quantified using *Dorado* (v7.2.0) with the “PseU basecaller”. In addition to the Ψ ratio provided by the basecaller, we calculated the U-to-C mismatch ratio, as previous studies have shown that Ψ incorporation may lead to U-to-C mismatches during base calling^42,43^, a feature not integrated in the “PseU basecaller” (**see supplementary notes S1**). To ensure selectivity, we filtered out sites with a modification ratio below 10% in each condition, and we used our IVT 18S as negative control.

Using this approach, we reproducibly identified all 104 Ψ sites in 18S, 5.8S, and 28S rRNAs previously identified by mass spectrometry^38^ (**Fig S10G, Source Data 3**). This demonstrate that NanoRibolyzer, coupled with “PseU basecaller”, is a robust tool for detecting ribosomal RNA pseudouridines. Using it in combination with C/U mismatch detection proved useful as it allowed to disregard hits at position not known to be modified, such as in the pre-rRNA spacers (**Fig S10G**, see e.g. green peaks in the 5’ ETS) Lastly, to gain information about spatio-temporal deposition of Ψ, we followed the ‘pre-rRNA sequential extraction’ (PSE) method^44^, to isolate cytoplasmic (Cp), nucleoplasmic (Np), and nucleolar (No) fractions, as confirmed by Western blot analysis with compartment-specific markers (**Fig 5A and S11A**).

**Figure 5.**
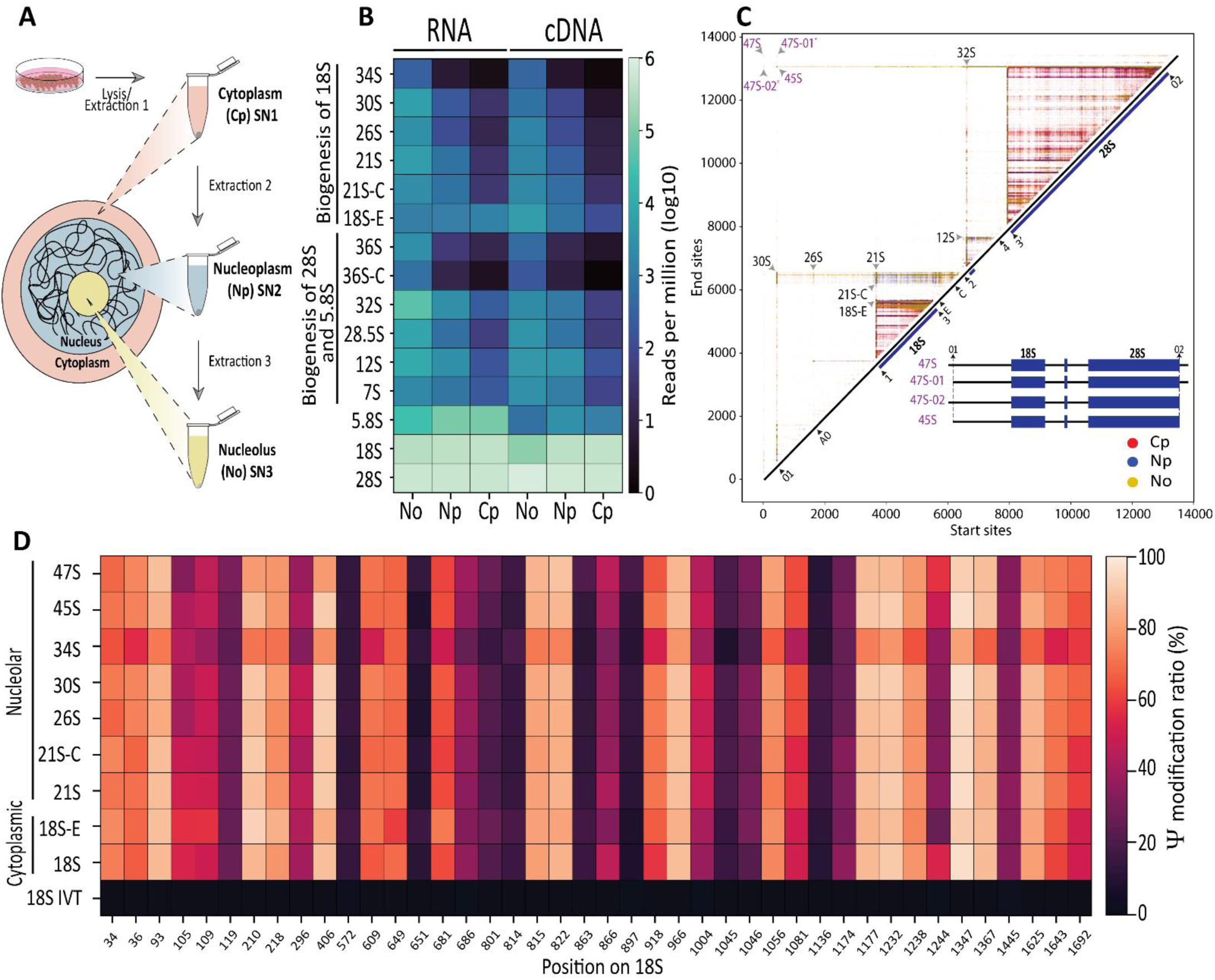
Spatio-temporal precursor-specific RNA modification detection and mapping of early precursors. **A**, Schematic diagram of pre-rRNA sequential extraction (PSE) method (Nieto et al. 2021). SN1, SN2 and SN3 represent the cytoplasmic (Cp), nucleoplasmic (Np) and nucleolar (No) fractions, respectively. **B**, Quantification of detected pre-rRNA intermediates and mature rRNAs in the nucleolus, nucleoplasm and cytoplasm (n=3) using direct RNA sequencing (RNA, left) and cDNA sequencing (right). **C**, Overlayed intensity matrix showing nucleolar (yellow), nucleoplasmic (blue), and cytoplasmic (red) samples. Key intensity hubs and their associated precursors are marked with arrows. The top left (purple) highlights the positions of 47S, 45S, and the 47S variants (47S-01 and 47S-02). The different rRNA products associated with 47S variants are illustrated in the center. **D**, Heatmap showing the pseudouridine modification ration across the 42 sites on 18S and classified by the associated precursors. The pseudouridine quantification was assessed according to precursors 47S, 45S, 30S, 26S, 21S, 21S-C and 18S-E which were quantified from the nucleolar fraction while the mature 18S was quantified in the cytoplasmic fraction. In-vitro transcribed 18S was used as negative control.

First, we performed cDNA sequencing in triplicate and analyzed coverage and precursor abundance across the distinct fractions (**Fig S11B-D)**. As expected, the nucleolar fraction (No) was highly enriched for transcripts spanning the 5’ and 3’ ETS and ITS1 and 2 regions (**Fig S11B)**. At the opposite range of the spectrum, transcripts detected in the cytoplasmic fraction (Cp) corresponded vastly to mature rRNAs. Transcripts enriched in the nucleoplasm (Np) had already undergone some level of maturation (**Fig S11B**).

Formal quantification revealed that the nucleolar fractions were enriched in the majority of pre-rRNA species, including 30S, 26S, 21S/21S-C, as well as 34S and 36S precursors – considered short-lived aberrant species^6^(**Fig S11C-D, Source Data 4**). Progressive processing was reflected by a gradient of precursors abundance, with the highest levels in the nucleolus, followed by the nucleoplasm and then the cytoplasm.

We next performed direct RNA sequencing (DRS), on these fractions to assess the dynamic deposition of Ψ from the nucleolus to the cytoplasm. Comparison of precursor abundance between cDNA and DRS sequencing were remarkably similar (**Fig 5B**, **Fig S11E-F**). Using an unsupervised approach, we identified in the DRS data, distinct processing hubs for key precursors, such as the previously detected 12S, 21S and 30S (see arrowheads in **Fig 5C**), and now also for early 45S and 47S in the nucleolar fraction (**Fig 5C and Fig S11G**). DRS offered higher resolution, lesser background, and captured longer precursors than cDNA sequencing. We believe this is due to differences in library preparation methods, and the use of newly released enzyme, which is more processive (**Fig S11G-H,** See **supplementary notes S2).**

Notably, we discovered novel early intermediates, which we tentatively named 47S-01 (lacking the sequence upstream of the 01 site) and 47S-02 (without the downstream sequence of the 02 site) (**Fig 5C**). These intermediates were already visible in the processing perturbation intensity matrices, particularly upon DIMT1L depletion (**Fig 4F**). We now have further proof by direct RNA sequencing they exist. These previously undocumented species presumably escaped detection due to the lower sensitivity and resolution of the methods available at the time.

Using the “PseU basecaller”, we reproducibly identified all the expected Ψ sites in fractions corresponding to the nucleolus, the nucleoplasm, and the cytoplasm (**Fig S12, Source Data 3)**. Focusing on the 42 Ψ’s found on 18S rRNA, for which we have unmodified *in vitro* synthesized control, and by performing precursor-specific mapping, we discovered that nearly all are already present on 47S, indicating cotranscriptional modification. We also noted that unfaithfully processed 34S species was less modified than “on path” precursors (**Fig 5D**). For a complete pseudouridylation frequency map, see **Supplementary Fig S13**. We conclude that most pseudouridines are already present in nucleolar RNAs. This provides, to our knowledge, the first demonstration that pseudouridylation is an early event.

In conclusion, NanoRibolyzer can resolve precursor-specific RNA modification dynamics, uncovering insights that were previously inaccessible due to technical limitations.

## Discussion

Ribosome biogenesis is a complex pathway involving hundreds of interconnected steps^1,2^. Among these, pre-rRNA processing (RNA cleavage) to generate mature rRNA ends serves as an excellent *proxy* for the overall process (**Fig 1A**). Traditionally, processing intermediates have been analyzed using metabolic labelling, northern blotting or primer extension. While these techniques are robust, they are somewhat limited in resolution, sensitivity, and throughput and often require access to costly and hazardous radioactive materials. This is particularly problematic for detecting low-abundance, metastable intermediates generated by progressive exoribonucleolytic trimming, such as the 3’ end maturation of the 18S and 5.8S rRNAs^7,8,9,10^(Fig 1A).

We have introduced NanoRibolyzer, an experimental and bioinformatic platform designed for the analysis and quantification of pre-rRNA intermediates using long-read Nanopore-sequencing. Coupled with a streamlined nuclei isolation protocol, we successfully detected both precursor and mature forms of nuclear and cytoplasmic rRNAs (**Fig 1B**). Utilizing long-read cDNA sequencing, which supports multiplexing for increased throughput and cost efficiency, we applied two complementary mapping approaches: a supervised (template-based) strategy for known precursors, and an unsupervised (template-free) approach, providing an agnostic characterization of reads across the entire 47S template (**Fig 2**).

The supervised approach enabled us to obtain quantitative measurements of rRNA precursors and mature rRNAs, which is particularly valuable for studying processing perturbations, such as the inactivation ribosome assembly factors (**Fig 4**).

Using the unsupervised approach, we detected pre-rRNAs associated with perturbations, defining processing fingerprints and identifying cleavage sites. Cross-referencing with the literature confirmed that these sites aligned with known processing sites, including 01(Ref.^23^), A0 (Ref.^24^), E (Refs ^25,26^), C (Ref.^25^) and 2 (Refs^27,28,29^) and others (**Fig 3**). While these sites exhibited sequence similarity at the nucleotide level, they did not always match the expected annotated positions. This highlights the importance of accurately documenting processing sites with nucleotide resolution using a standard reference sequence. We have summarized our findings on cleavage site definition in **Fig 6**.

**Figure 6.**
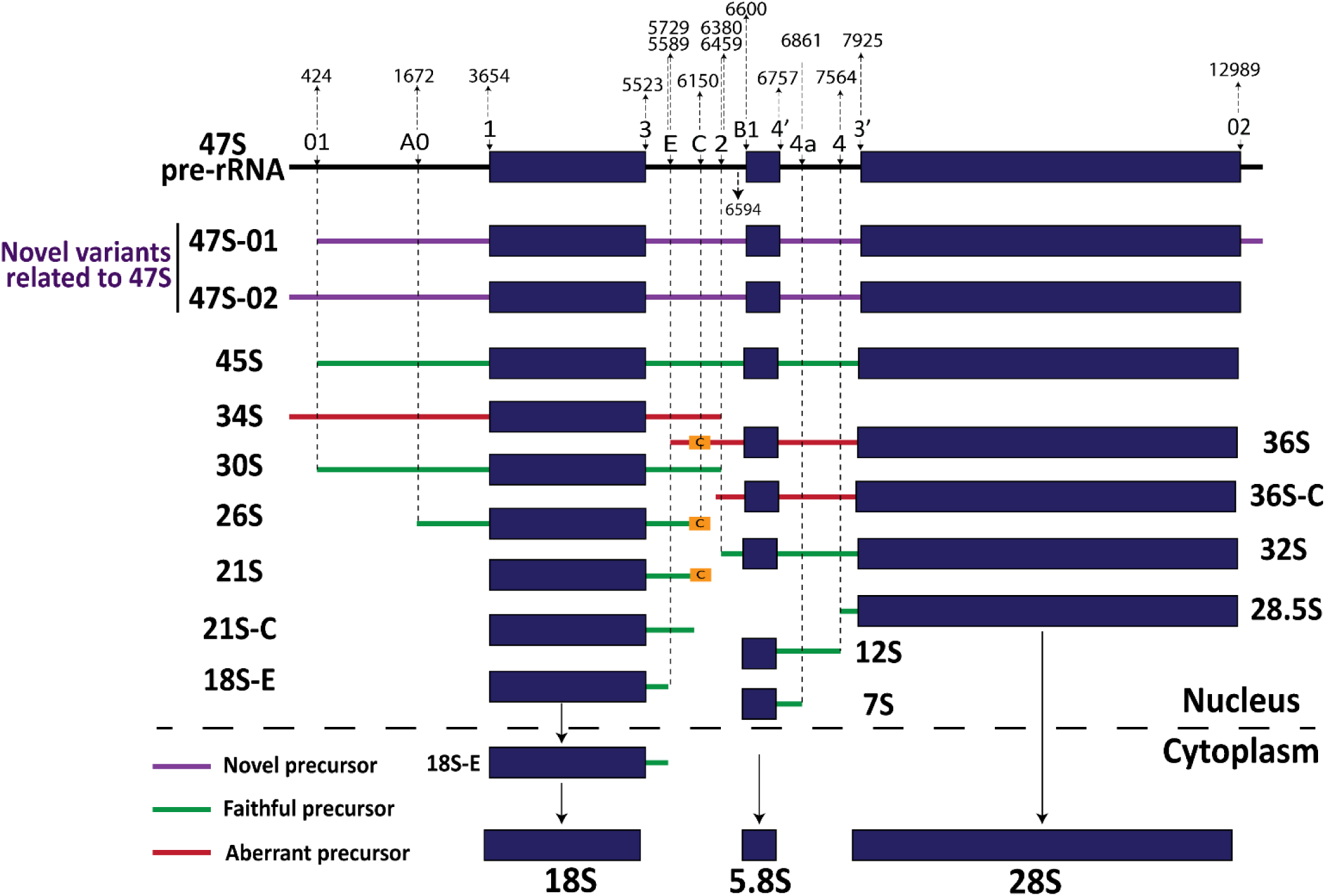
Overview of redefined processing sites across the human 47S pre-rRNA. Refined nucleotide-level annotations of all processing sites are shown along the top of the 47S pre-rRNA. Through combined nuclear fractionation, and both supervised and unsupervised analytical approaches, we identified previously unreported 47S intermediates, likely due to the limited resolution of traditional methods such as northern blotting.

Additionally, we identified novel putative processing sites and alternative precursors, providing new insights into human ribosome biogenesis. For instance, we confirmed two potential precursors associated with site 2: one corresponding to the previously described 21S precursor (site 2^1^, Ref. ^27,28^) and a shorter 21S variant (site 2^2^,), which had been observed in a previous study^29^. We also predicted the putative location of site 4, which had not been fully characterized. In the 2-D matrix display, we observed what we refer to as processing “smears” indicating exoribonucleolytic digestion within spacers regions. We believe these correspond to degradation resulting from aborted subunit production, surveillance, and/or stress-induced activation of damage pathways^4,45,46^. These observations suggest that NanoRibolyzer could also serve as a powerful tool for exploring rRNA surveillance and turnover.

NanoRibolyzer not only confirmed the effects of known processing perturbations but also provided higher-resolution insights into their impact. For example, we found that depletion of the small subunit factor DIMT1L leads to accumulation of the 3′ ETS and 32S –an observation not previously reported (**Fig 4D**). We also showed that RPL3 depletion, like URB1, resulted in altered processing at the 3′ ETS. Additionally, for those two factors, we observed altered read coverage at processing site 01, suggesting potential coupling between 5′ and 3′ end maturation (**Fig 4C**). These findings highlight NanoRibolyzer’s potential in detecting processing defects and revealing the distinct molecular “fingerprints” associated with each perturbation.

Using DRS, we presented a proof-of-concept for spatial RNA modification analysis, enabling the detection of RNA modifications in both precursor and mature rRNA (**Fig 5**). Our template-based approach classified pre-rRNA precursors and using raw signal data, profiled the dynamic spatial states of three base modifications, including m⁷G, 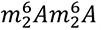, and m¹acp³Ψ. Using a pseudouridine-specific basecaller in combination with C/U mismatch detection, we uncovered that pseudouridines were already present in 47S precursor (**Fig 6**), indicating co-transcriptional deposition. While cotranscriptional modification has previously been shown for 2′-O-methylation in yeast^47^, our findings provide evidence that pseudouridylation also occurs cotranscriptionally in human cells. In contrast, the aberrant 34S precursor was less modified, suggesting RNA modification may play a role in controlling “on path” ribosome biogenesis, as suggested for other base modifications^34,48^.

In summary, NanoRibolyzer provides a simple and adaptable nanopore-based approach to study pre-rRNA processing and modification in human cells, overcoming limitations of previous complex or narrowly focused methods. By enabling single-nucleotide resolution and quantitative analysis of pre-rRNA intermediates, it offers valuable insights into ribosome biogenesis, with broad applications in basic research, disease understanding (ribosomopathies), and clinical diagnostics.

## Material and methods

### Cell lines and culture

HEK293 cells (ATCC CRL-1573) were cultured in DMEM supplemented with 10% FBS, and 1% L-glutamine and maintained in an incubator at 37°C and 5% CO_2_.

All mutations analyses were introduced in homozygous diploid in HCT116 p53 positive cells.

| Cell line | Ref. |
| --- | --- |
| **tsr3 -/-** | 39 |
| **dimt1l-Y131G** | 49 |
| **wbsr22-D82K** | 49 |

### siRNA inactivation experiments

Cells were revere transfected with silencers (10 nM, except URB1: 15 nM) in a time course (H6, H12, H24, H48, H72) to identify the best condition for Nanoribolyzer analysis^6^. All DsiRNAs silencers were used at 10 nM final (except for URB1, 15 nM). Silencers against UTP18, WBSCR22, DIMT1L, and LAS1L are Silencer Select (Ambion). Silencers against URB1 and RPL3 are DsiRNAs (IDT).

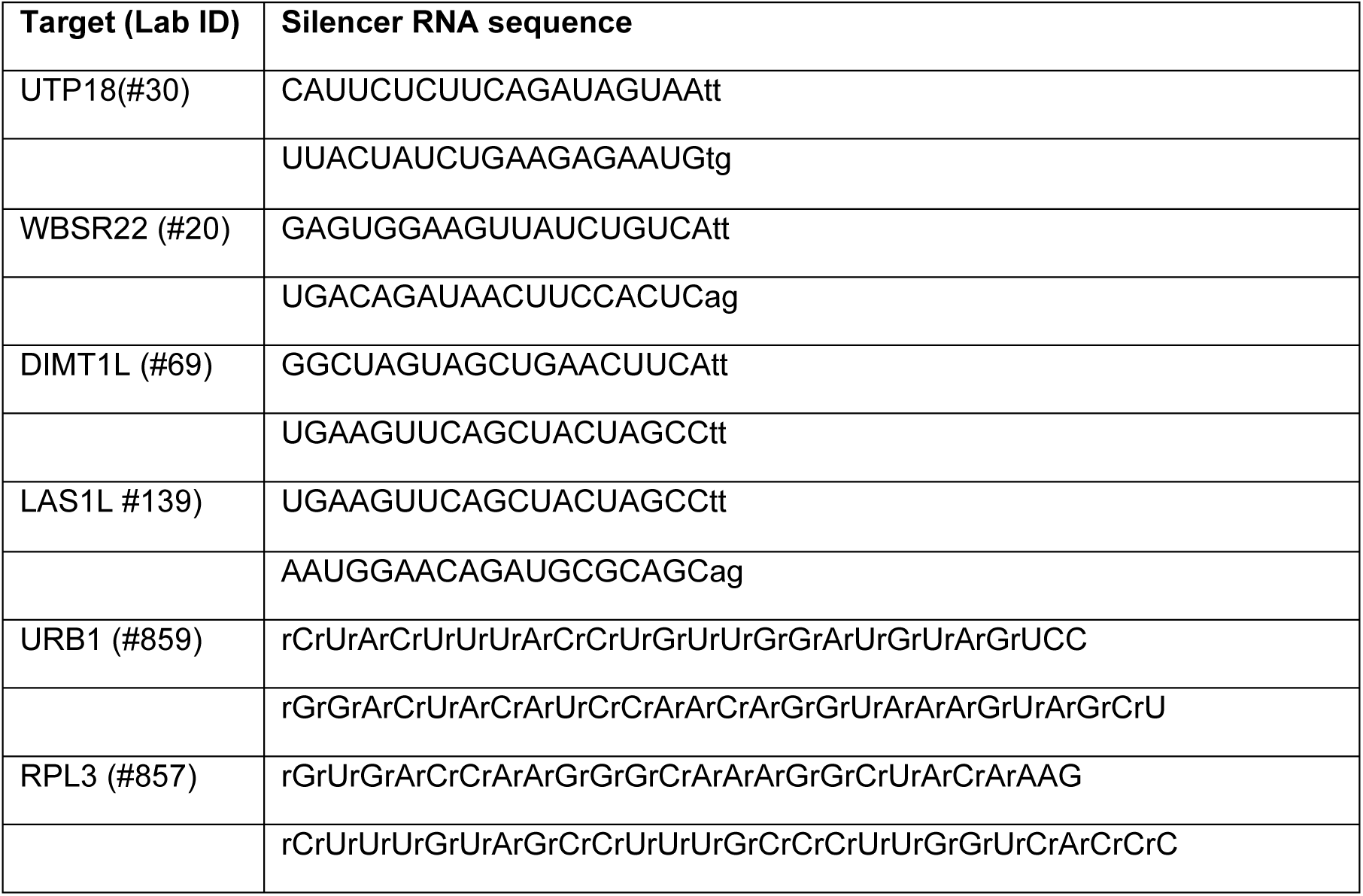

### Simplified nuclei isolation protocol

Detailed description of the protocol is illustrated in the Supplementary Fig S1. Briefly, samples were trypsined and washed with cold PBS, after which they were resuspended in Nuclei Isolation Buffer (NIB - 10mM Tris-HCl (pH7.4),10mM NaCl, 3mM MgCl2, 0.1% Igepal, 0.1% Tween-20,1% BSA, 0.15mM Spermine, 0.15mM Spermidine, 0.2U/ul RNase inhibitor) and homogenized using a loose pestle with ten strokes. The homogenate was then incubated on ice for 15 minutes and subsequently centrifuged at 800 rpm for 5 minutes at 4 °C. The resulting soluble fraction (cytoplasmic) was transferred to a new tube, and Trizol was added to the sample at room temperature (RT) while nuclei isolation continued. The remaining pellet was subjected to two washes with 500 μl of ice-cold NIB buffer and centrifuged at 800rpm for 5 minutes at 4 °C. After the second wash, the supernatant was removed, and the pellet was resuspended with 300 μl of ice-cold NIB buffer. For nuclei isolation an equal volume (300 μl) of 50% Optiprep solution (Stemcell technologies, 07820) was added to the homogenate sample and thoroughly resuspended by pipetting, resulting in a 25% sample/optiprep mix. Then, 600 μl of 40% Optiprep solution, followed by 30% Optiprep solution, were layered in a 2 ml Eppendorf tube. The 25% sample/optiprep mix was layered on top of the 30% solution forming three visible layers. The tube was centrifuged at top speed (14,000rpm) in a bench centrifuge for 20 minutes at 4 °C. After centrifugation, the nuclei (∼600 μl) were carefully collected from the 40%-30% phase and transferred into a 1.5 ml Lo-bind tube. 600 μl of Nuclei Wash Buffer (NWB - 10mM Tris-HCl (pH7.4),10mM NaCl, 3mM MgCl2, 0.1% Tween-20,1% BSA, 0.2U/ul RNase inhibitor) was added to the nuclei and thoroughly resuspended by pipetting. The sample was then centrifuged at 1,100rpm for 10 minutes at 4 °C, and the supernatant was removed. The nuclei were washed again with 500 μl of ice-cold NWB buffer and centrifuged similarly. The supernatant was removed, leaving approximately 20 μl of solution. Quality of the nuclei was assessed using DAPI staining and visualized under the microscope using a UV filter to identify DAPI-positive nuclei. High-quality nuclei were characterized by debris-free, round or oval-shaped DAPI-stained nuclei. Once the quality of isolated nuclei was confirmed, remaining nuclei samples were treated with DNaseI (NEB). The DNase I master mix was supplemented with 0.15 mM spermidine, 0.15 mM spermine, and RNAse inhibitor (U/ul) per sample. The samples were then incubated at 37 °C for 20 minutes. Next, the sample volume was brought to 500 μl using nuclease-free water, and 500 μl of TRIzol reagent (Life Technologies, 15596026) was added to the nuclei samples to proceed with RNA isolation.

### Pre-ribosomal sequential extraction method (PSE)

The PSE method was performed as previously described^44^. Briefly, cells (HEK293) were grown at at ∼80% confluency and harvested on ice-cold phosphate-buffered saline. Cell were resupended thoroughly in 0.5 ml of SN1 buffer (20 mM HEPES-NaOH [pH 7.5], 130 mM KCl, 10 mM MgCl2, 0.05% Igepal CA-630), supplemented with Cømplete protease inhibitor cocktail (Roche), followed by centrifugation 1300 × g for 3 min at 4 °C. The supernatant was collected and stored as the SN1 fraction. The pellet was washed with 0.5 ml SN1 buffer, and then resuspended in 0.3 ml of SN2 buffer (10 mM HEPES-NaOH (pH 7.5), 10 mM NaCl, 5 mM MgCl2, 0.1% Igepal CA-630, 0.5 mg/ml heparin, 600 U/ml RNasin (Promega)) supplemented with 100 U RNase-free DNase I (Qiagen), and incubated for 10 min at room temperature with gentle mixing. The lysate was centrifuged 12,300 × g for 10 min at 4 °C and the supernatant collected as the SN2 fraction. The remaining pellet was resuspended in 0.4 ml of SN3 buffer (20 mM HEPES-NaOH (pH 7.5), 200 mM NaCl, 4 mM EDTA, 0.1% Igepal CA- 630, 0.04% sodium deoxycholate, 4 mM imidazole, 0.1 mg/ml heparin, 1 mM dithiothreitol (DTT), Cømplete, 600 U/ml RNasin) and incubated for 20 min at room temperature with moderate agitation. The extract was centrifuged 12,300 × g for 10 min in 4 °C and the supernatant was collected as the SN3 fraction. Total RNA was prepared from each fraction using TRIzol reagent (Life Technologies, 15596026) according to the manufacturer’s protocol.

### RNA isolation

RNA isolation was performed as previously described^50^. Nuclear and cytoplasmic (as well as whole cell) fractions (∼1 mL) were incubated in RT for at least 5 Min. 200µl Chlorophorm was added to nuclear and cytoplasmic fractions. Samples were Vortexed for 15 sec, incubated in RT fort 3 min and centrifuged for 15 min full speed (FS) at 4 °C. The upper aqueous phase (∼550 µl) was transferred into a fresh Eppendorf tube and 500 µl isopropanol was added, thoroughly resuspended and incubated in RT for 15 min. Sample were centrifuge 10 min at full speed at 4 °C. The supernatant was discarded pellet was washed with 75% EtOH in nuclease free water followed by a 5 min centrifugation at 7500g at 4°C. The supernatant was discarded, and pellet was air dried for 5 – 10 min. RNA was eluted with 30 µl of RNAse free water and mix in Hula mixer for 10 min in RT. RNA concentration was measured with qubit and RNA integrity was assessed via Agilent RNA ScreenTape analysis.

### Northern Blotting

Total RNA was extracted using TRIzol reagent, separated on agarose denaturing gel, and analyzed by northern blotting, as previously described^6^. The depletion optimization was performed on HEK293 cells where each depletion factor displayed the expected phenotype according to the literature. On the basis of this analysis, the following depletion time points were selected for NanoRibolyzer: UTP18 (48h), DIMT1L (72h), WBSCR22 (48h), LAS1L (48h), and URB1 (72h) (see Fig 3b).

#### Probes for northern blotting

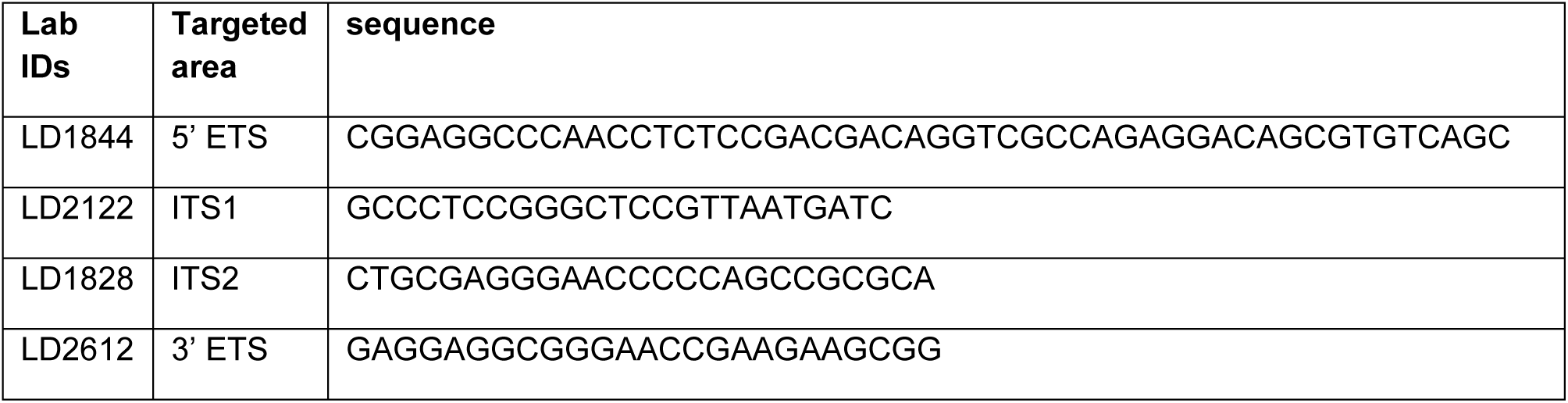

### Western blot controls for SN1, SN2, and SN3 fractionation

Equivalent amounts of SN1, SN2, and SN3 fractions, each extracted from starting material equivalent to 25 µg of total protein (fractionated according to^44^), along with 25 µg of total protein from whole-cell lysate used as control, were separated on a 10% SDS-PAGE gel. Proteins were transferred to nitrocellulose membranes and probed in TBS containing 0.1% Tween-20 and 3% BSA. Membranes were incubated with either: 1) Anti-rabbit primary antibodies: anti-histone H3 (Proteintech, Ref 17168- 1-AP), anti-nucleolin (Proteintech, Ref 10556-1-AP), and anti-α-tubulin (Proteintech, Ref 11224-1-AP), each at 1:5000 for 2 hours, followed by HRP-conjugated anti-rabbit secondary antibody (Cytiva, Ref NA934V) at 1:5000 for 2 hours, or 2) Anti-mouse primary antibodies: anti-fibrillarin (Antibodies-online, Ref ABIN361375) or anti-UBF (Tebu Bio, Ref SC-13125), each at 1:1000 overnight, followed by HRP-conjugated anti-mouse secondary antibody (Jackson ImmunoResearch, Ref 115-035-062) at 1:5000 for 2 hours.

### In-vitro poly adenylation using poly(A) tailing of RNA

For in-vitro polyadenylation, “In-vitro poly adenylation using poly(A) tailing of RNA” kit was used (NEB#M0276), according to the manufacturer instructions. Briefly, 1 µg of RNA was taken in 15 µl nuclease free water and supplemented with 2 µl of 10X E. coli Poly(A) Polymerase Reaction Buffer, 2 µl ATP (10mM) and 1 µl E. coli Poly(A) Polymerase (to a total volume 20 µl). Samples were incubated at 37°C for 30 minutes. The polyadenylated RNA samples were purified using RNAClean XP beads (Beckman Coulter, A63987), according to the manufacturer instructions. In the last elution step, the sample was resuspended in 10 µl nuclease-free water and incubated at 37°C for 5 minutes in Hula mixer. The sample was placed on the magnet and once the solution was clear, the elute was transferred into a clean 1.5 ml Eppendorf tube.

### Direct cDNA-native barcoding library preparation

Direct cDNA coupled with native barcoding libraries were prepared using Direct cDNA Sequencing Kit (SQK-DCS109), Native Barcoding Expansion 1-24 (EXP-NBD104, EXP-NBD114), following the manufacturer’s protocol. *Reverse transcription and strand-switching.* 1µg of poly(A)-tailed RNA was transferred to a 1.5 ml tube and adjusted to 7.5 µl with nuclease-free water. In a 0.2 ml PCR tube, 7.5 µl of RNA sample were mixed with 2.5 µl of VNP (ONT), 2.5 µl of 10 mM dNTPs (NEB N0447), and the volume was adjusted to 11 µl with nuclease-free water. The samples were incubated for 10 minutes at room temperature and then snap-cooled on a pre-chilled freezer block for 1 minute. Next, a master mix was prepared, containing 4 μl of 5x RT Buffer (ThermoFisher, EP0751), 1 μl RNaseOUT (Life Technologies, 10777019), 1 μl of Nuclease-free water, and 2 μl Strand-Switching Primer (SSP, ONT) per sample, to a total volume of 8 μl. The strand-switching buffer was added to the snap-cooled, annealed mRNA, and the samples were incubated at 42°C for 2 minutes in the thermal cycler. Subsequently, 1 µl of Maxima H Minus Reverse Transcriptase (ThermoFisher, EP0751) was added, and the total volume becomes 20 µl. The samples were incubated following a specific thermal protocol: 42°C for 90 mins, 85°C for 5 mins, and then holding at 4°C. After the reverse transcription, RNA degradation and second strand synthesis were performed. 1 µl of RNase Cocktail Enzyme Mix (ThermoFisher, AM2286) was added to the reverse transcription reaction and incubated for 10 minutes at 37°C. The samples were then subjected to the AMPure XP beads-based (Beckman Coulter A63881) purification method using 0.85x ratio of beads:sample and ultimately cDNA hybrid was eluted in 20 µl of nuclease-free water. Next, the 20 μl of reverse-transcribed samples were prepared with 25 μl of 2x LongAmp Taq Master Mix (NEB, N0447), 2 μl of PR2 Primer (PR2, ONT), and 3 μl of Nuclease-free water, to a total volume of 50 μl. The samples were incubated at specific temperatures in the thermocycler. Afterwards, the sample were then subjected to the AMPure XP beads-based purification method using 0.8x ratio of beads:sample and ultimately cDNA/RNA hybrid was eluted in 21 µl of nuclease-free water. The eluted sample was quantified using a Qubit fluorometer. *End-prep.* Subsequently, end repair and dA-tailing were performed by mixing 20 µl of cDNA sample with 30 µl Nuclease-free water, 7 µl Ultra II End-prep reaction buffer (NEB, E7546), and 3 µl Ultra II End-prep enzyme (NEB, E7546) mix to a total volume of 60 µl. The samples were incubated at 20°C for 5 minutes and then at 65°C for 5 minutes. Next, the samples were subjected to AMPure XP beads-based purification using 1x ratio of beads:sample, and the cDNA was eluted with 22.5 µl of nuclease-free water. *Barcode ligation.* Barcode ligation was then performed, where 22.5 µl of End-prepped DNA was mixed with 2.5 µl of Native Barcode and 25 µl of Blunt/TA Ligase Master Mix (NEB, M0367) to a total volume of 50 µl. The reaction was incubated for 15 minutes at room temperature. The samples were then subjected to AMPure XP beads-based purification using 1x ratio of beads:sample, and the cDNA was eluted with 26 µl of nuclease-free water. Lastly, the barcoded samples are pooled to a final volume of 65 μl in a 1.5 ml Eppendorf tube. *Adapter ligation.* Adapter ligation was performed by adding 65 µl of pooled barcoded sample, 5 µl of Adapter Mix II (AMII, ONT), 20 µl of 5X NEBNext Quick Ligation Reaction Buffer (NEB, B6058), and 10 µl of Quick T4 DNA Ligase (NEB, E6056) to a total volume of 100 µl. The final libraries were incubated for 10 minutes at room temperature and then subjected to AMPure XP beads-based purification, with the cDNA being eluted with 26 µl of nuclease-free water. The sample was loaded and sequenced onto a primed PromethION flow cell as per the manufacturer instruction.

### Direct RNA library preparation

Direct RNA libraries were prepared using the SQK-RNA004 kit (ONT) following the manufacturer’s protocol. Briefly, 1µg of poly(A)-tailed RNA was adjusted to final volume of 9.5 µl with nuclease-free water. 3 µl of NEBNext Quick Ligation Reaction Buffer (NEB B6058), 1 µl of RT Adapter (RTA) (ONT), and 1.5 µl of T4 DNA Ligase 2M U/ml (NEB M0202), were added to sample resulting in a total volume of 15 µl. The reaction is mixed by pipetting and incubated for 10 minutes at room temperature. Next, the reverse transcription master mix was prepared by mixing 9 µl of Nuclease-free water, 2 µl of 10 mM dNTPs (NEB N0447), 8 µl of 5x first-strand buffer (Thermo Fisher Scientific, 18080044), and 4 µl of 0.1 M DTT, resulting in a total volume of 23 µl. The master mix was added to the RNA sample containing the RT adapter-ligated RNA. 2 µl of SuperScript III reverse transcriptase (Thermo Fisher Scientific, 18080044) were added to the reaction, bringing the final volume to 40 µl. The reaction was incubated at 50°C for 50 minutes, followed by 70°C for 10 minutes, and then brought to 4°C. Agencourt RNAClean XP beads (Beckman Coulter, A63987) were resuspended and 72 µl of the resuspended beads were added to the reaction. The sample was mixed by pipetting and incubated on a Hula mixer for 5 minutes at room temperature. Subsequently, the sample was subjected to two washes with 70% ethanol, and the RNA:DNA hybrids were eluted with 20 µl of nuclease-free water. For the adapter ligation reaction, 8.0 µl of NEBNext Quick Ligation Reaction Buffer, 6.0 µl of RNA Ligation Adapter (RLA), 3.0 µl of Nuclease-free water, and 3.0 µl of T4 DNA Ligase were mixed with 20 µl of the eluted sample to reach a total volume of 40 µl. The reaction was incubated for 10 minutes at room temperature. 30 µl of resuspended RNAClean XP beads were added to the adapter ligation reaction, mixed by pipetting, and incubated on a Hula mixer for 5 minutes at room temperature. The sample was then subjected to two washes with the Wash Buffer (WSB, ONT) using a magnetic rack. Following the washes, the beads were pelleted on the magnet, and the supernatant was pipetted off. The pellet was resuspended in 33 µl of Elution Buffer (EB, ONT) and were incubated at 37°C for 10 minutes in a Hula mixer. Incubation at 37°C allows the release of long fragments from the beads. The eluate was then cleared by pelleting the beads on a magnet, and the eluate was retained and transferred to a clean to 1.5 ml tube. The sample was loaded and sequenced onto primed PromethION flow cell as per the manufacturer instruction.

### Implementation of NanoRibolyzer pipeline

NanoRibolyzer was implemented as a Nextflow-based workflow using Docker containers and could be installed as a plugin within Oxford Nanopore Technologies’ (ONT) Epi2Me platform (https://github.com/stegiopast/wf-nanoribolyzer). Pod5 output files were basecalled using the dorado basecaller (https://github.com/nanoporetech/dorado), trimmed with Porechop (https://github.com/rrwick/Porechop) to remove adapter sequences and aligned to the 45SN1 (equivalent to 47S) (GeneID:106631777; NW_021160023.1:480347-493697) using minimap2^51^ with the map-ont flag. The read IDs of the aligned 45SN1 were used to filter the original pod5 file. The resulting unaligned BAM files were used to collect metadata on reads at a single nucleotide resolution and the rebasecalled reads were realigned to the 45SN1 reference. The final BAM files were processed using both template-based and template-free approaches.

NanoRibolyzer is open-source and has been made freely available to the community through the Epi2ME platform for broader accessibility and use.

### Template-based quantification of rRNA precursors

The template-based algorithm associated long-reads with literature-based ribosomal intermediates^8,9,10^. In this approach, the pairwise minimal reciprocal overlap (MRO) between a query read and all possible intermediates was determined. The MRO was defined by calculating the minimal relative overlap of the query over the intermediate and vice versa. Once the minimal overlap for each query-intermediate pair was established, the pair with the maximal overlap was used to associate the read with the corresponding intermediate (See Fig S3). Read clusters were then stored in a tab-separated values (TSV) table, which included read IDs, absolute and relative read counts, and the start and end sites of all reads associated with each intermediate. Additionally, bed files for each intermediate were generated to facilitate visualization in the Integrative Genomics Viewer^52^ (IGV). The 45SN1 reference FASTA from the NanoRibolyzer references repository was used for all analyses.

### Template-free rRNA precursors

The template-free algorithm was based on the construction of a 2-dimensional (length(45SN1)^2^) intensity matrix in which reads were embedded using the alignment start and end sites as coordinates. The number of reads sharing start and end site coordinates on the matrix led to the formation of intensity “hubs”. The resulting intensity matrix was stored in CSV format, which included the start site, end site, number of reads, and ID list for each intensity hub. For visualization of the matrices, absolute read counts of intensity hubs were min-max normalized applying an additional contrast enhancement of 2%.

### Determination of read cleavage sites

The determination of significant abundant cleavage sites was computed using alignment start and end sites where the absolute abundance of start and end sites was determined along the 45SN1 reference. For each template-based intermediate, the mean abundance and standard deviation (SD) of alignment start and end sites were calculated. Cleavage sites occurring two SDs above the mean in the dataset were considered significant abundant cleavage sites. For overlapping template-based intermediate cleavage site intervals, metrics were determined by calculating the mean of means and the mean of SDs. The output files were stored in TSV and BED file formats, including information about their relative abundance and the cleavage site location. The BED files were visualized using IGV.

### DRS Modification analysis

PseudoU detection was performed by using dorado version 7.2.0 with the super accuracy RNA004 model version 5.0.0 using the “pseU” flag. The resulting unaligned bam file was converted into fastq format using samtools^53^ (https://github.com/samtools/samtools). The fastq file was aligned with minimap2^51^ using the “map-ont” long read alignment flag together with the MD flag to ensure the remainder of the modification information. Subsequently, the aligned bam file was processed using pysam (https://github.com/pysam-developers/pysam). For each possibly modified nucleotide on a read aligned bam files contain a modification probability. A threshold of 95% modification probability was chosen to determine modifications.

For the acquisition of the raw signal analysis at specific modification sites, the data was basecalled and aligned as described above. Template-based analysis information was used to determine intermediate-specific datasets. The aligned bam file and the original pod5 together with a 45SN1 reference fasta file were processed using remora (https://github.com/nanoporetech/remora). Remora was used to perform the resquiggleing, which is the process of association of raw signal interval with specific nucleotides on the reference or basecalled sequence. In this case, once the intervals were associated to the reference sequence, the mean of z-normalized raw current signal covering 10 bases up- and downstream from the loci of interest were extracted. For all reads covering the locus of interest the mean of means and semi-standard deviation (deviation below and above mean) of means were determined.

### Statistics

All statistical analyses are described in the respective figure legends. Each legend provides detailed information about the statistical metrics (such as the mean and standard deviation), sample sizes, statistical tests used and any relevant adjustments applied.

## Code availability

All scripts and code used in this work have been made available on GitHub (https://github.com/stegiopast/wf-nanoribolyzer).

## Acknowledgments

This work was partly funded by Deutsche Forschungsgemeinschaft (DFG, German Research Foundation; project no. 439669440 TRR319 RMaP TP A05/C01/C03 to M.H. and S.M). T.B and S.G. acknowledge funding from the ReALity initiative of the Johannes Gutenberg University Mainz. Research in the Lab of D.L.J.L. was supported by the Belgian Fonds de la Recherche Scientifique (F.R.S./FNRS), EOS [CD-INFLADIS, grant n°40007512], Région Wallonne (SPW EER) Win4SpinOff [RIBOGENESIS], the COST actions EPITRAN (CA16120) and TRANSLACORE (CA21154), the European Joint Programme on Rare Diseases (EJP-RD) RiboEurope and DBAGeneCure. S.G. and L.L acknowledge funding by the Boehringer Ingelheim Stiftung. S.G. acknowledge funding by SFB 1552 Project No. 465145163 of the Deutsche Forschungsgemeinschaft (DFG).

## Contributions

S.P. developed and implemented the NanoRibolyzer pipeline; S.P. and L.L. analyzed the data. T.B. L.W. and S.P. performed nuclei isolation and RNA isolation experiments. L.W. performed the knockdown experiments and northern blots. S.M. performed the in-vitro transcription of 18S. T.B. and S.P. prepared and sequenced the Nanopore-seq libraries. T.B. and D.L. conceived and supervised the work, with the assistance of S.G., M.H. and B.L., which provided valuable input and feedback in various discussions. T.B., D.L. and S.P. wrote the paper, with contributions from all the authors.

## Ethics declarations

All authors declare that the research was conducted in the absence of any commercial or financial relationships that could be construed as a potential conflict of interest.

## References

1 Lafontaine, D. L. J. (2015). Noncoding RNAs in eukaryotic ribosome biogenesis and function. Nature Structural & Molecular Biology, 22(1), 11–19. 10.1038/NSMB.2939

2 Bohnsack, K. E., & Bohnsack, M. T. (2019). Uncovering the assembly pathway of human ribosomes and its emerging links to disease. The EMBO Journal, 38(13). 10.15252/EMBJ.2018100278

3 Ni, C., & Buszczak, M. (2023). Ribosome biogenesis and function in development and disease. Development (Cambridge, England), 150(5). 10.1242/DEV.201187

4 Kang, J., Brajanovski, N., Chan, K. T., Xuan, J., Pearson, R. B., & Sanij, E. (2021). Ribosomal proteins and human diseases: molecular mechanisms and targeted therapy. Signal Transduction and Targeted Therapy 2021 6:1, 6(1), 1–22. 10.1038/s41392-021-00728-8

5 Popov, A., Smirnov, E., Kováčik, L., Raška, O., Hagen, G., Stixová, L., & Raška, I. (2013). Duration of the first steps of the human rRNA processing. Nucleus, 4(2), 134–141. 10.4161/NUCL.23985

6 Tafforeau, L., Zorbas, C., Langhendries, J. L., Mullineux, S. T., Stamatopoulou, V., Mullier, R., Wacheul, L., & Lafontaine, D. L. J. (2013). The complexity of human ribosome biogenesis revealed by systematic nucleolar screening of Pre-rRNA processing factors. Molecular Cell, 51(4), 539–551. 10.1016/J.MOLCEL.2013.08.011

7 Grummt, I. (2013). The nucleolus—guardian of cellular homeostasis and genome integrity. Chromosoma, 122(6), 487–497. 10.1007/S00412-013-0430-0

8 Tomecki, R., Sikorski, P. J., & Zakrzewska-Placzek, M. (2017). Comparison of preribosomal RNA processing pathways in yeast, plant and human cells – focus on coordinated action of endo- and exoribonucleases. FEBS Letters, 591(13), 1801–1850. 10.1002/1873-3468.12682

9 Henras, A. K., Plisson-Chastang, C., O’Donohue, M. F., Chakraborty, A., & Gleizes, P. E. (2015). An overview of pre-ribosomal RNA processing in eukaryotes. Wiley Interdisciplinary Reviews: RNA, 6(2), 225–242. 10.1002/WRNA.1269

10 Mullineux, S. T., & Lafontaine, D. L. J. (2012). Mapping the cleavage sites on mammalian pre-rRNAs: where do we stand? Biochimie, 94(7), 1521–1532. 10.1016/J.BIOCHI.2012.02.001

11 Hall, A. N., Morton, E., & Queitsch, C. (2022). First discovered, long out of sight, finally visible: ribosomal DNA. Trends in Genetics : TIG, 38(6), 587–597. 10.1016/J.TIG.2022.02.005

12 McStay, B. (2016). Nucleolar organizer regions: genomic “dark matter” requiring illumination. Genes & Development, 30(14), 1598–1610. 10.1101/GAD.283838.116

13 Agrawal, S., & Ganley, A. R. D. (2018). The conservation landscape of the human ribosomal RNA gene repeats. PLOS ONE, 13(12), e0207531. 10.1371/JOURNAL.PONE.0207531

14 Floutsakou, I., Agrawal, S., Nguyen, T. T., Seoighe, C., Ganley, A. R. D., & McStay, B. (2013). The shared genomic architecture of human nucleolar organizer regions. Genome Research, 23(12), 2003–2012. 10.1101/GR.157941.113

15 Wang, Y., Zhao, Y., Bollas, A., Wang, Y., & Au, K. F. (2021). Nanopore sequencing technology, bioinformatics and applications. Nature Biotechnology 2021 39:11, 39(11), 1348–1365. 10.1038/s41587-021-01108-x

16 Kono, N., & Arakawa, K. (2019). Nanopore sequencing: Review of potential applications in functional genomics. Development, Growth & Differentiation, 61(5), 316–326. 10.1111/DGD.12608

17 Amarasinghe, S. L., Su, S., Dong, X., Zappia, L., Ritchie, M. E., & Gouil, Q. (2020). Opportunities and challenges in long-read sequencing data analysis. Genome Biology, 21(1). 10.1186/S13059-020-1935-5

18 Zheng, P., Zhou, C., Ding, Y., Liu, B., Lu, L., Zhu, F., & Duan, S. (2023). Nanopore sequencing technology and its applications. MedComm, 4(4), e316. 10.1002/MCO2.316

19 Butto, T., Mungikar, K., Baumann, P., Winter, J., Lutz, B., & Gerber, S. (2023). Nuclei on the Rise: When Nuclei-Based Methods Meet Next-Generation Sequencing. Cells 2023, Vol. 12, Page 1051, 12(7), 1051. 10.3390/CELLS12071051

20 Begik, O., Diensthuber, G., Liu, H., Delgado-Tejedor, A., Kontur, C., Niazi, A. M., Valen, E., Giraldez, A. J., Beaudoin, J. D., Mattick, J. S., & Novoa, E. M. (2023). Nano3P-seq: transcriptome-wide analysis of gene expression and tail dynamics using end-capture nanopore cDNA sequencing. Nature Methods, 20(1), 75–85. 10.1038/S41592-022-01714-W

21 Brand, R.C., Klootwijk, J., Planta, R.J. and Maden, B.E. (1978)Biosynthesis of a hypermodified nucleotide in Saccharomycescarlsbergensis 17S and HeLa-cell 18S ribosomal ribonucleic acid.Biochem J., 169, 71–77

22 Pirouz, M., Munafò, M., Ebrahimi, A. G., Choe, J., & Gregory, R. I. (2019). Exonuclease requirements for mammalian ribosomal RNA biogenesis and surveillance. Nature Structural & Molecular Biology 2019 26:6, 26(6), 490–500. 10.1038/s41594-019-0234-x

23 Kass, S., Craig, N., & Sollner Webbi, B. (1987). Primary Processing of Mammalian rRNA Involves Two Adjacent Cleavages and Is Not Species Specific. MOLECULAR AND CELLULAR BIOLOGY, 7(8), 2891–2898.

24 Rouquette, J., Choesmel, V., & Gleizes, P. E. (2005). Nuclear export and cytoplasmic processing of precursors to the 40S ribosomal subunits in mammalian cells. The EMBO Journal, 24(16), 2862–2872. 10.1038/SJ.EMBOJ.7600752

25 Preti, M., O’Donohue, M. F., Montel-Lehry, N., Bortolin-Cavaillé, M. L., Choesmel, V., & Gleizes, P. E. (2013). Gradual processing of the ITS1 from the nucleolus to the cytoplasm during synthesis of the human 18S rRNA. Nucleic Acids Research, 41(8), 4709. 10.1093/NAR/GKT160

26 Montellese, C., Montel-Lehry, N., Henras, A. K., Kutay, U., Gleizes, P. E., & O’Donohue, M. F. (2017). Poly(A)-specific ribonuclease is a nuclear ribosome biogenesis factor involved in human 18S rRNA maturation. Nucleic Acids Research, 45(11), 6822–6836. 10.1093/NAR/GKX253

27 Goldfarb, K. C., & Cech, T. R. (2017). Targeted CRISPR disruption reveals a role for RNase MRP RNA in human preribosomal RNA processing. Genes and Development, 31(1), 59–71. 10.1101/GAD.286963.116/-/DC1

28 Idol, R. A., Robledo, S., Du, H.-Y., Crimmins, D. L., Wilson, D. B., Ladenson, J. H., Bessler, M., & Mason, P. J. (2007). Cells depleted for RPS19, a protein associated with Diamond Blackfan Anemia, show defects in 18S ribosomal RNA synthesis and small ribosomal subunit production. 10.1016/j.bcmd.2007.02.001

29 Ishikawa, H., Yoshikawa, H., Izumikawa, K., Miura, Y., Taoka, M., Nobe, Y., Yamauchi, Y., Nakayama, H., Simpson, R. J., Isobe, T., & Takahashi, N. (2017). Poly(A)-specific ribonuclease regulates the processing of small-subunit rRNAs in human cells. Nucleic Acids Research, 45(6), 3437–3447. 10.1093/NAR/GKW1047

30 Schillewaert, S., Wacheul, L., Lhomme, F., & Lafontaine, D. L. J. (2012). The Evolutionarily Conserved Protein LAS1 Is Required for Pre-rRNA Processing at Both Ends of ITS2. Molecular and Cellular Biology, 32(2), 430. 10.1128/MCB.06019-11

31 Sloan, K. E., Mattijssen, S., Lebaron, S., Tollervey, D., Pruijn, G. J. M., & Watkins, N. J. (2013). Both endonucleolytic and exonucleolytic cleavage mediate ITS1 removal during human ribosomal RNA processing. Journal of Cell Biology, 200(5), 577–588. 10.1083/JCB.201207131

32 Hölzel, M., Orban, M., Hochstatter, J., Rohrmoser, M., Harasim, T., Malamoussi, A., Kremmer, E., Längst, G., & Eick, D. (2010). Defects in 18 S or 28 S rRNA processing activate the p53 pathway. The Journal of Biological Chemistry, 285(9), 6364–6370. 10.1074/JBC.M109.054734

33 Haag, S., Kretschmer, J., & Bohnsack, M. T. (2015). WBSCR22/Merm1 is required for late nuclear pre-ribosomal RNA processing and mediates N7-methylation of G1639 in human 18S rRNA. RNA (New York, N.Y.), 21(2), 180–187. 10.1261/RNA.047910.114

34 Zorbas, C., Nicolas, E., Wacheul, L., Huvelle, E., Heurgué-Hamard, V., & Lafontaine, D. L. J. (2015). The human 18S rRNA base methyltransferases DIMT1L and WBSCR22-TRMT112 but not rRNA modification are required for ribosome biogenesis. Molecular Biology of the Cell, 26(11), 2080– 2095. 10.1091/MBC.E15-02-0073

35 Castle, C. D., Cassimere, E. K., Lee, J., & Denicourt, C. (2010). Las1L is a nucleolar protein required for cell proliferation and ribosome biogenesis. Molecular and Cellular Biology, 30(18), 4404– 4414. 10.1128/MCB.00358-10

36 Shan, L., Xu, G., Yao, R. W., Luan, P. F., Huang, Y., Zhang, P. H., Pan, Y. H., Zhang, L., Gao, X., Li, Y., Cao, S. M., Gao, S. X., Yang, Z. H., Li, S., Yang, L. Z., Wang, Y., Wong, C. C. L., Yu, L., Li, J., … Chen, L. L. (2023). Nucleolar URB1 ensures 3′ ETS rRNA removal to prevent exosome surveillance. Nature 2023 615:7952, 615(7952), 526–534. 10.1038/s41586-023-05767-5

37. de La Cruz, J., Karbstein, K., & Woolford, J. L. (2015). Functions of ribosomal proteins in assembly of eukaryotic ribosomes in vivo. Annual Review of Biochemistry, 84, 93–129. 10.1146/ANNUREV-BIOCHEM-060614-033917

38 Taoka, M., Nobe, Y., Yamaki, Y., Sato, K., Ishikawa, H., Izumikawa, K., Yamauchi, Y., Hirota, K., Nakayama, H., Takahashi, N., & Isobe, T. (2018). Landscape of the complete RNA chemical modifications in the human 80S ribosome. Nucleic Acids Research, 46(18), 9289–9298. 10.1093/NAR/GKY811

39 Meyer, B., Wurm, J. P., Sharma, S., Immer, C., Pogoryelov, D., Kötter, P., Lafontaine, D. L. J., Wöhnert, J., & Entian, K. D. (2016). Ribosome biogenesis factor Tsr3 is the aminocarboxypropyl transferase responsible for 18S rRNA hypermodification in yeast and humans. Nucleic Acids Research, 44(9), 4304–4316. 10.1093/NAR/GKW244

40 Barozzi, C., Zacchini, F., Asghar, S., & Montanaro, L. (2022). Ribosomal RNA Pseudouridylation: Will Newly Available Methods Finally Define the Contribution of This Modification to Human Ribosome Plasticity? Frontiers in Genetics, 13, 920987. 10.3389/FGENE.2022.920987/BIBTEX

41 Pryszcz, L. P., Diensthuber, G., Llovera, L., Medina, R., Delgado-Tejedor, A., Cozzuto, L., Ponomarenko, J., & Novoa, E. M. (2025). Rapid and accurate demultiplexing of direct RNA nanopore sequencing data with SeqTagger. Genome Research, 35(4), 956–966. 10.1101/GR.279290.124/-/DC1

42 Hewel, C., Hofmann, F., Dietrich, V., Wierczeiko, A., Friedrich, J., Jenson, K., Mündnich, S., Diederich, S., Sys, S., Schartel, L., Schweiger, S., Helm, M., Lemke, E. A., Linke, M., & Gerber, S. (2024). Direct RNA sequencing (RNA004) allows for improved transcriptome assessment and near real-time tracking of methylation for medical applications. BioRxiv, 2024.07.25.605188. 10.1101/2024.07.25.605188

43 Schartel, L., Jann, C., Wierczeiko, A., Butto, T., Mündnich, S., Marchand, V., Motorin, Y., Helm, M., Gerber, S., & Lemke, E. A. (2024). Selective RNA pseudouridinylation in situ by circular gRNAs in designer organelles. Nature Communications 2024 15:1, 15(1), 1–10. 10.1038/s41467-024-53403-1

44 Nieto, B., Gaspar, S. G., Moriggi, G., Pestov, D. G., Bustelo, X. R., & Dosil, M. (2020). Identification of distinct maturation steps involved in human 40S ribosomal subunit biosynthesis. Nature Communications 2020 11:1, 11(1), 1–17. 10.1038/s41467-019-13990-w

45 King, K. L., Jewell, C. M., Bortner, C. D., & Cidlowski, J. A. (2000). 28S ribosome degradation in lymphoid cell apoptosis: evidence for caspase and Bcl-2-dependent and -independent pathways. Cell Death & Differentiation 2000 7:10, 7(10), 994–1001. 10.1038/sj.cdd.4400731

46 Parker, M. D., & Karbstein, K. (2023). Quality control ensures fidelity in ribosome assembly and cellular health. Journal of Cell Biology, 222(4). 10.1083/JCB.202209115/213871

47 Koš, M., & Tollervey, D. (2010). Yeast Pre-rRNA Processing and Modification Occur Cotranscriptionally. Molecular Cell, 37(6), 809–820. 10.1016/j.molcel.2010.02.024

48 Lafontaine, D., Vandenhaute, J., & Tollervey, D. (1995). The 18S rRNA dimethylase Dim1p is required for pre-ribosomal RNA processing in yeast. Genes and Development, 9(20), 2470–2481. 10.1101/GAD.9.20.2470

49. Naarmann-de Vries, I. S., Zorbas, C., Lemsara, A., Piechotta, M., Ernst, F. G. M., Wacheul, L., Lafontaine, D. L. J., & Dieterich, C. (2023). Comprehensive identification of diverse ribosomal RNA modifications by targeted nanopore direct RNA sequencing and JACUSA2. RNA Biology, 20(1), 652– 665. 10.1080/15476286.2023.2248752

50 Butto, T., Pastore, S., Müller, M., Iyer, K. V., Mündnich, S., Wierczeiko, A., Friedland, K., Helm, M., Winz, M.-L., & Gerber, S. (2024). Real-time transcriptomic profiling in distinct experimental conditions. ELife, 13. 10.7554/ELIFE.98768.1

51 Li, H. (2018). Minimap2: pairwise alignment for nucleotide sequences. Bioinformatics, 34(18), 3094–3100. 10.1093/BIOINFORMATICS/BTY191

52 Robinson, J. T., Thorvaldsdóttir, H., Winckler, W., Guttman, M., Lander, E. S., Getz, G., & Mesirov, J. P. (2011). Integrative genomics viewer. Nature Biotechnology, 29(1), 24–26. 10.1038/NBT.1754

53 Danecek, P., Bonfield, J. K., Liddle, J., Marshall, J., Ohan, V., Pollard, M. O., Whitwham, A., Keane, T., McCarthy, S. A., & Davies, R. M. (2021). Twelve years of SAMtools and BCFtools. GigaScience, 10(2). 10.1093/GIGASCIENCE/GIAB008

