## Supplementary Material for "Mapping human pre-rRNA processing and modification at single nucleotide resolution using long read Nanopore sequencing"

### Supplementary Figures

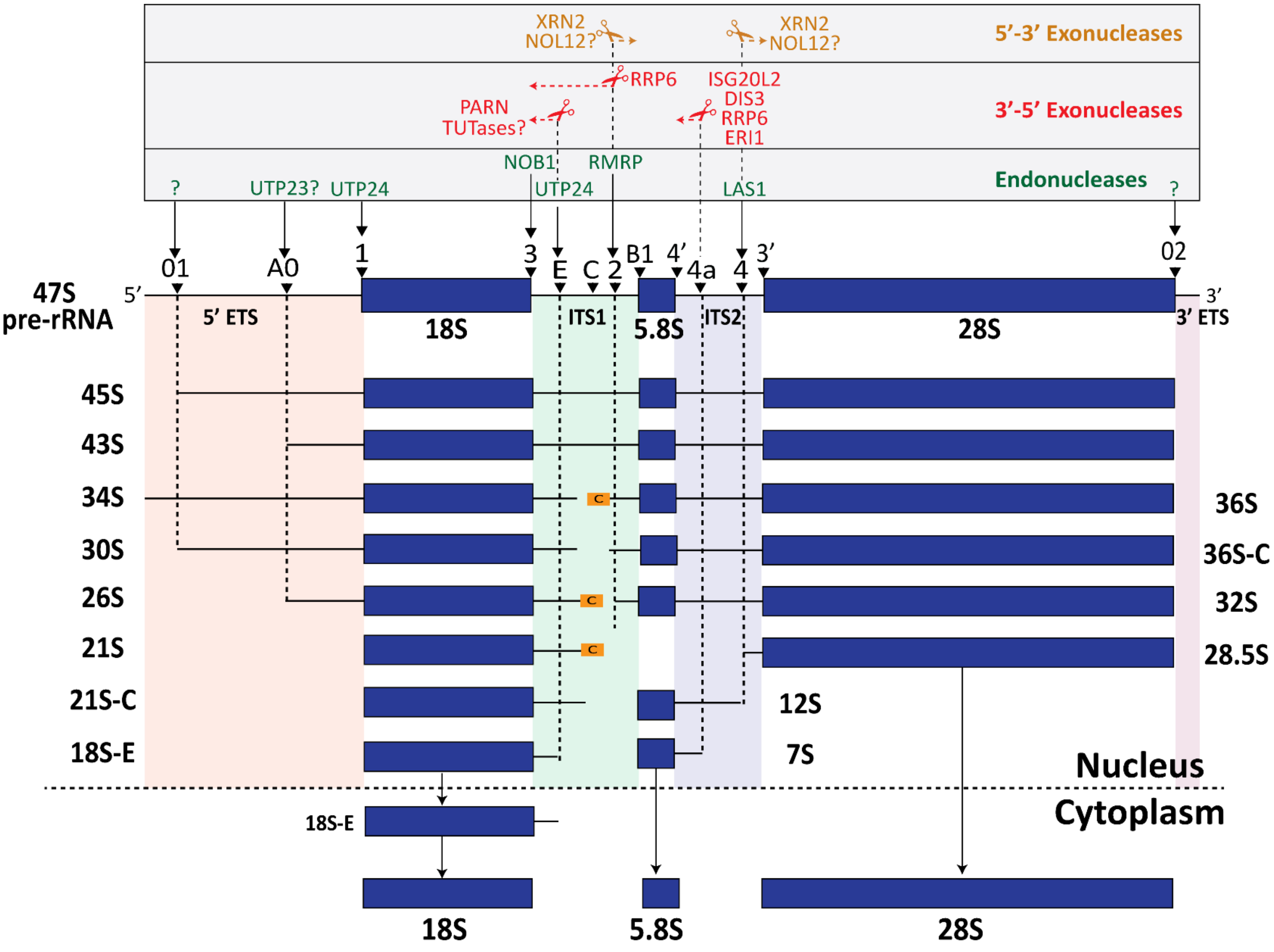


***Supplementary Figure S1. Overview of human pre-rRNA processing pathway.***
Schematic overview of processing sites in the human 47S pre-rRNA, summarizing over two decades of research on rRNA processing within the external transcribed spacers (5′ ETS and 3′ ETS) and internal transcribed spacers (ITS1 and ITS2), shown below the 47S template. Enzymes involved in specific processing steps are indicated: endonucleases (green), 3′-5′ exonucleases (red), 5′-3′ exonucleases (orange), and unknown enzymes (?).
The boundaries for the mature rRNA components are indicated as follows: 18S rRNA (sites 1 and 3), 5.8S rRNA (sites B1 and 4'), and 28S rRNA (sites 3' and 02).
Early precursors include 47S, 45S and 43S. The precursors for 18S rRNA biogenesis include 30S, 26S, 21S, 21S-C, and 18S-E, while the precursors for 28S and 5.8S rRNA include 32S, 28.5S, 12S, and 7S [8,9,10]. The 36S, 36S-C, and 34S RNAs were associated with dysfunctional subunit biogenesis.

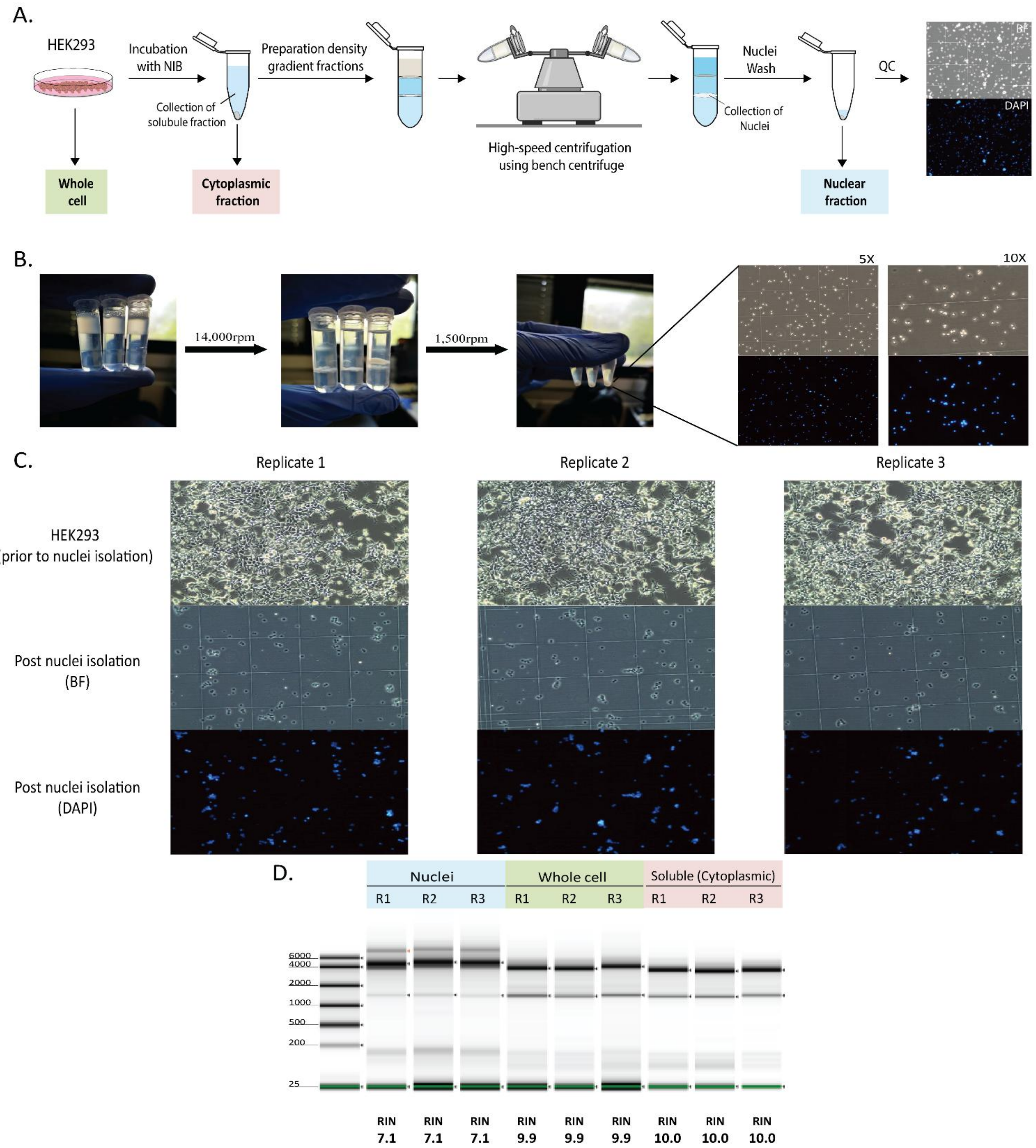


***Supplementary Figure S2. Simplified nuclei isolation procedure and quality control steps.***
**A,** Detailed schematic diagram of the simplified nuclei isolation procedure. See materials and methods for details. Cells are incubated in nuclei isolation buffer (NIB) for 15 min. Following low speed centrifugation, the soluble fraction (containing the cytoplasmic fraction) is collected and stored for Trizol RNA isolation (light red). Density gradient solutions are prepared from 60% optiprep solution forming three layers where the pellet containing nuclei are placed at the top. Following high-speed centrifugation using a bench centrifuge, the nuclei are collected, washed and assessed under a microscope. Nuclei are then stored in Trizol for RNA isolation (light blue). The whole cell condition is collected at the first step and stored for Trizol RNA isolation as control (light green).
**B,** Illustration of the 2 ml tubes with density gradient and sample at the top of the tube. Following high-speed centrifugation (~14k rpm), the lower interphase (containing the nuclei) is collected and washed using low-speed centrifugation (~1.5k rpm). A sample from pelleted nuclei is taken, incubated with DAPI, and analyzed under bright field microscopy with UV filter to visualize the quality of the nuclei.

**C**, Representative microscope image of HEK293 cells (top) prior nuclei isolation, bright field post nuclei isolation (middle) and DAPI stained nuclei post nuclei isolation (bottom).
**D**, Tape-station analysis of RNA profiles of cytoplasmic (light red), whole cell (light green) and nuclear (blue) samples (n=3). RIN, RNA integrity number.

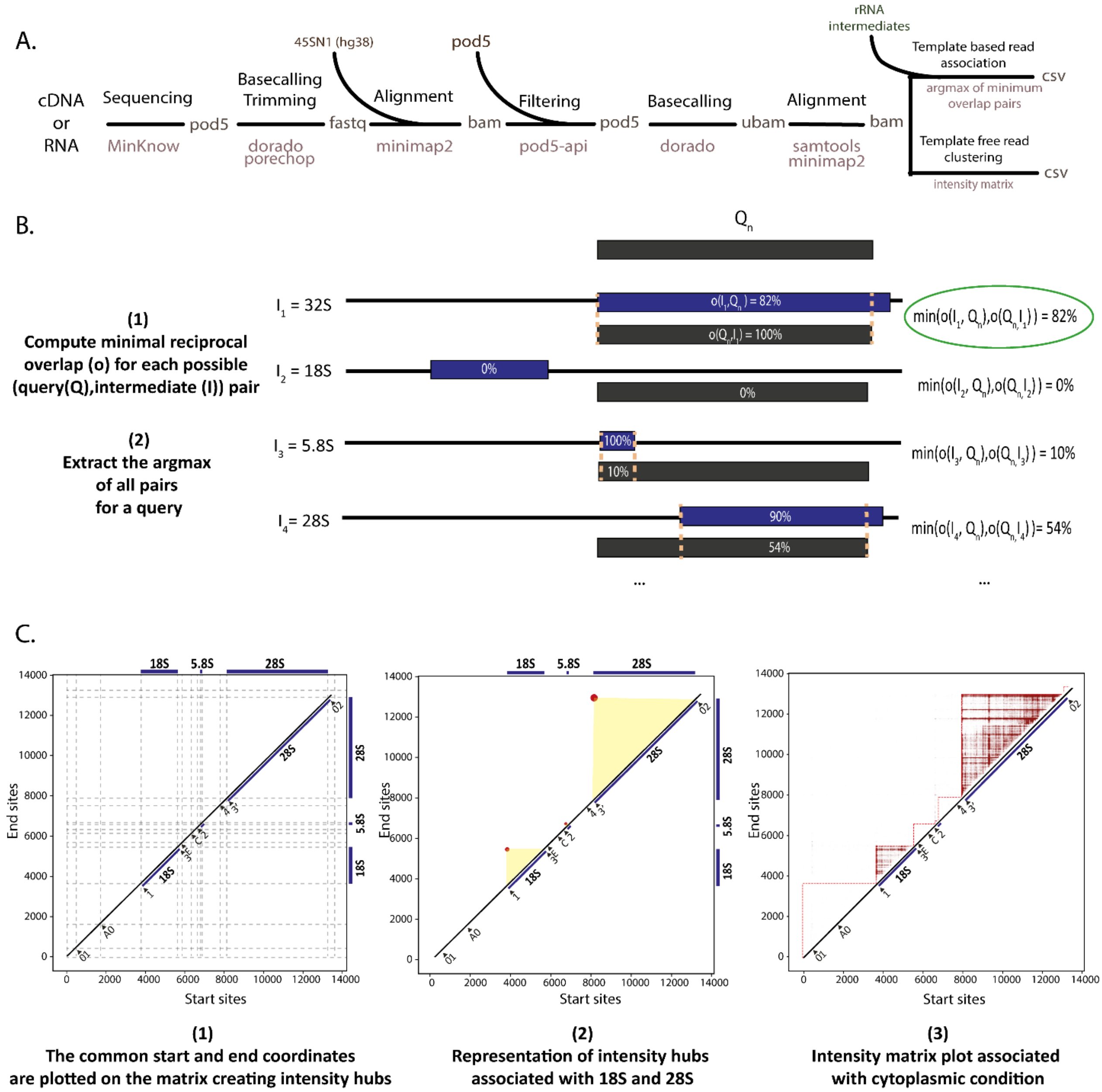


***Supplementary Figure S3. Overview of the NanoRibolyzer bioinformatic pipeline.***

**A,** NanoRibolyzer pipeline. Reads of input pod5 become basecalled and trimmed. Alignment to 45SN1 of hg38 is used to extract reads aligning to ribosomal RNA. Extracted reads become rebasecalled to perform polyA-estimation and modification detection. Rebasecalled reads are realigned to 45SN1 of hg38. Template-free and template-based read association analysis is performed on realigned reads.

**B,** Supervised or template-based approach. Minimal reciprocal overlap (MRO) between query read and all literature-based intermediates is computed based on alignment start and end sites of the query. The query read is associated with the intermediate with the maximal query/intermediate MRO. Formal: I={Intermediates}, Q={Query reads}, A={Associated query reads}, Qn ∈ Q , $A_n$=argmax(min(overlap($I_i$,$Q_n$),overlap($Q_n$,$I_i$)) ∀ $I_i$ ∈ I).

**C,** unsupervised or template-free approach. A 2-D matrix at the length of the RNA45SN1 template is constructed. Augmentation of reads by start and end site coordinate according to the alignment reveals intensity hubs. Intensity hubs are min-max normalized with a contrast enhancement of 2%.

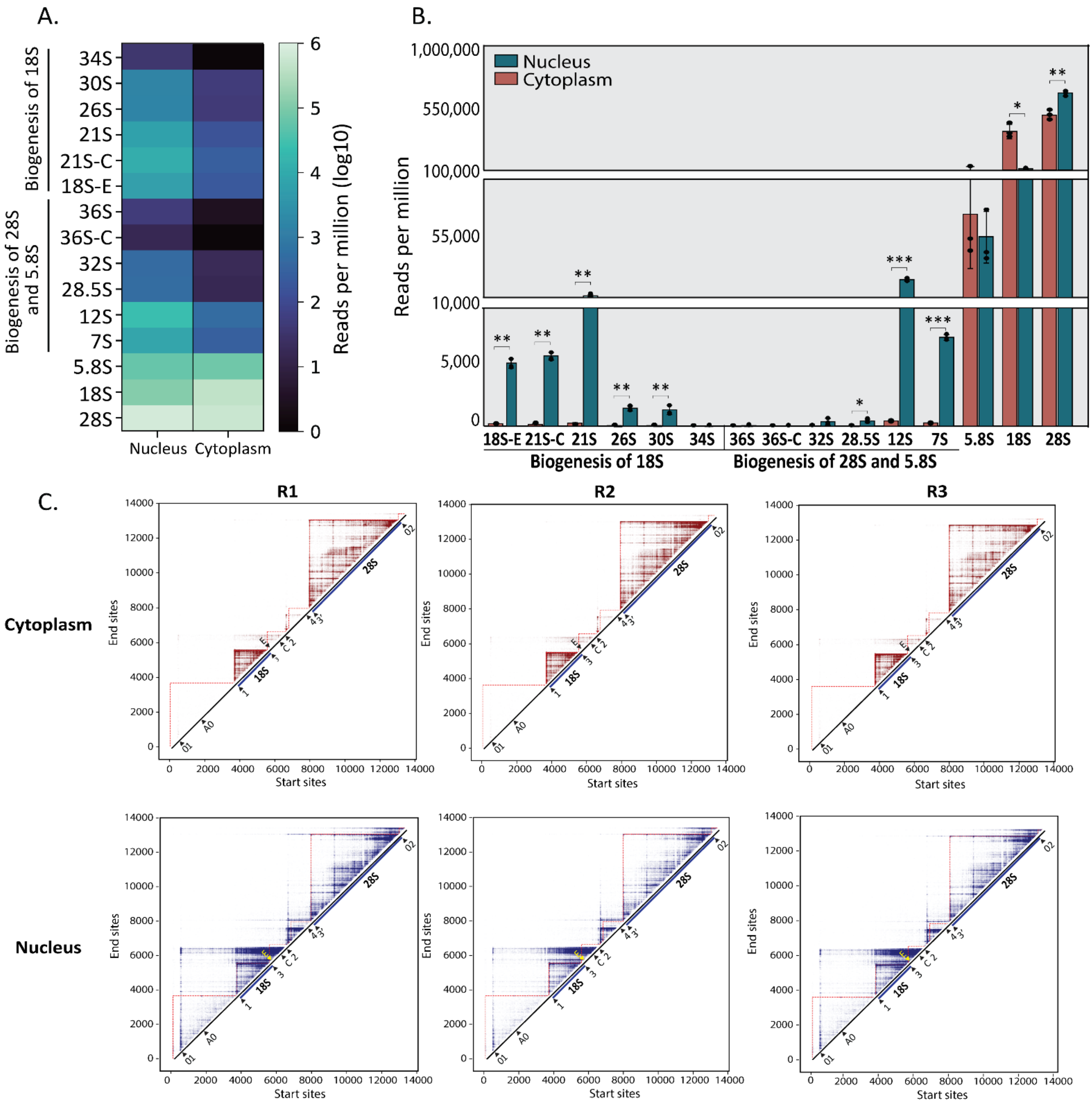


***Supplementary Figure S4. Pre-rRNA intermediate visualization and quantification in nucleus, cell and cytoplasm.***

**A,** Heatmap illustrating pre-/rRNA abundance in nuclear and cytoplasmic fractions, represented as log10 reads per million (averaged across three replicates).

**B**, Quantitative comparison of pre-rRNA intermediates and mature rRNA between nuclear and cytoplasmic fractions (n=3 each). Histograms display mean reads per million ± SD. Statistical significance was determined using a t-test between nucleus and cytoplasm conditions, with $*p < 0.05$, $**p < 0.01$, $***p < 0.001$.

**C**, Intensity matrices of cytoplasm and nuclei conditions (n=3).

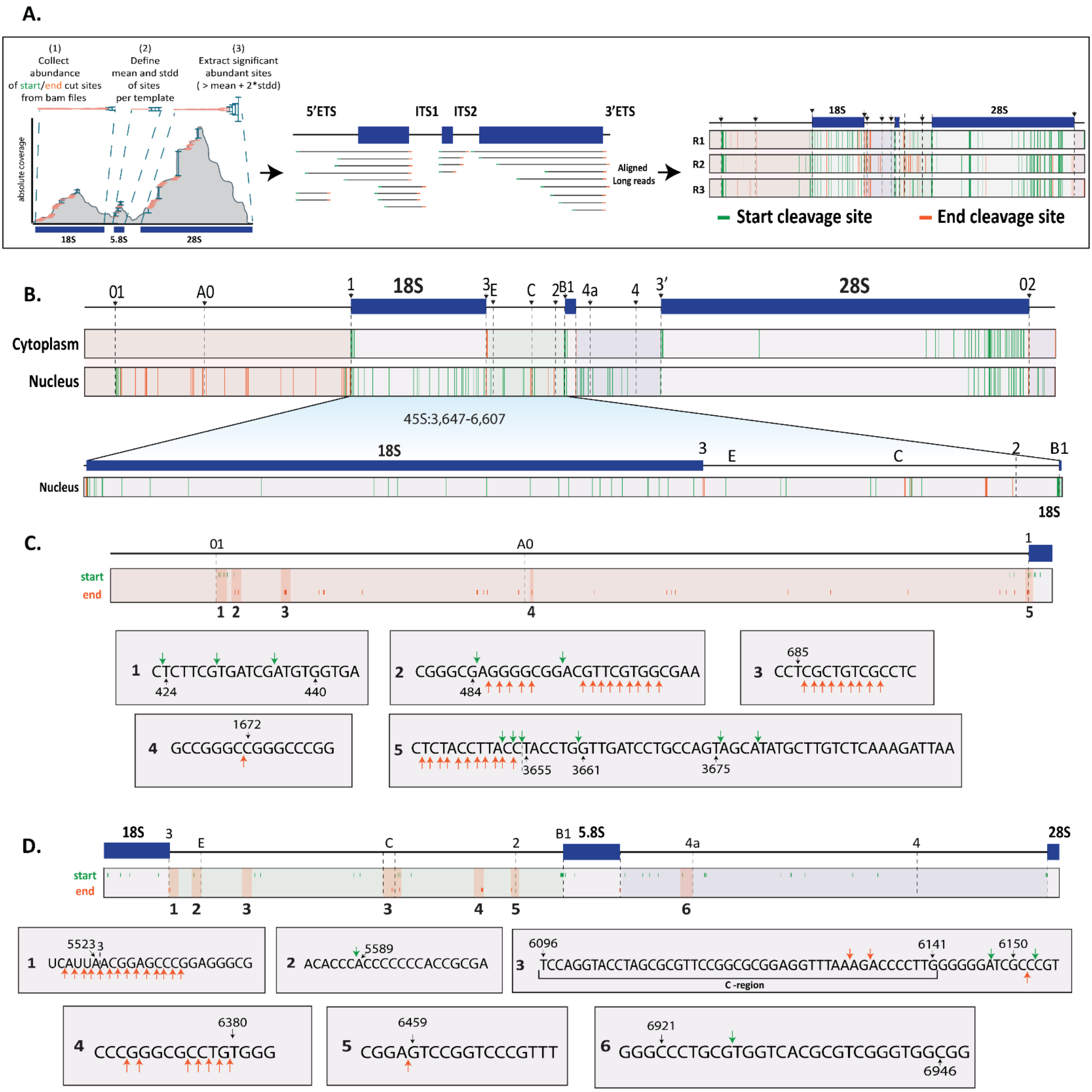


***Supplementary Figure S5. Determination and visualization of cleavages sites following processing perturbations.***

**A,** Extraction of prevalent cleavage sites strategy. Prevalent cleavage sites are extracted from the template-based precursors by collecting their start (green) and end (orange) sites for each precursor. The mean and standard deviation of these sites are calculated, and significant cleavage sites are those with abundances at least two standard deviations above the mean. The results are then saved in TSV and BED formats for visualization.

**B,** Visual representation of prevalent start (green) and end (orange) cleavage sites across the 47S rRNA in cytoplasm (top) and nucleus (bottom). Close-up view of prevalent cleavage sites at the 18S and ITS1 regions are shown below. Putative processing sites and mature rRNA sequences are annotated at the top of each respective figure. The coordinates spanning the region on the 47S rRNA are displayed above.

**C-D,** Detailed annotation of cleavage sites at single nucleotide resolution. Selected sites near bona fide processing sites within the 5' ETS (C), ITS1 and ITS2 (D), are highlighted, numbered, and the corresponding numbers are mapped to the actual sequence. The annotated sequences include nucleotide positions (black arrows), as well as start (green arrows) and end (orange arrows) cleavage sites, derived from cDNA data. Each cleavage site displayed was detected in at least two out of three replicates.

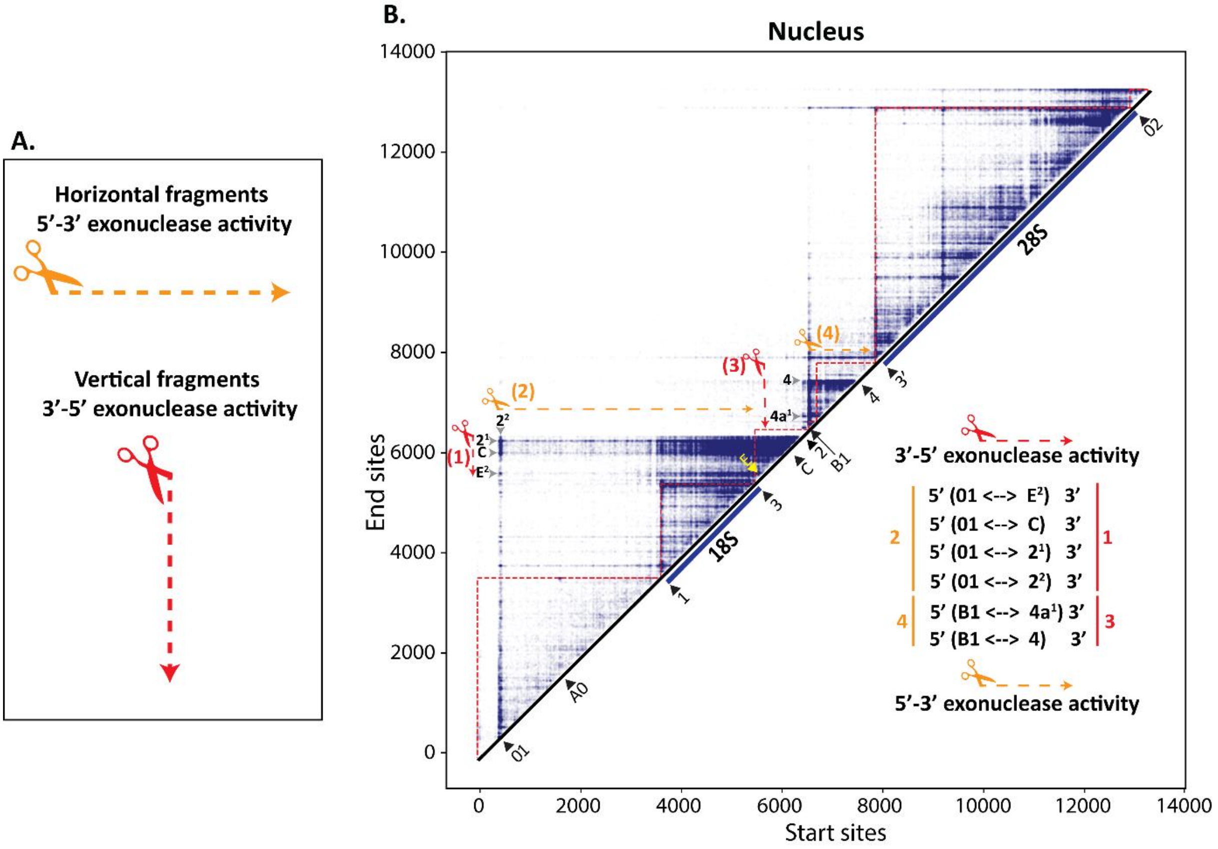


***Supplementary Figure S6. Exonucleolytic cleavage identification using the intensity matrix.***

**A,** The intensity matrix provides a clear visualization of exonucleolytic cleavage patterns: Horizontal fragmentation indicates a retained end processing site, with altered start sites, consistent with 5′-3′ exonuclease activity (orange). Vertical fragmentation indicates a retained start processing site, with altered end sites, consistent with 3′-5′ exonuclease activity (red).

**B**, Retained end processing sites are marked in black within the intensity matrix in the nuclear fraction. The 5’ and 3’ processing sites are indicated on the right with the directionality of exonuclease acitvity colored appropriately.

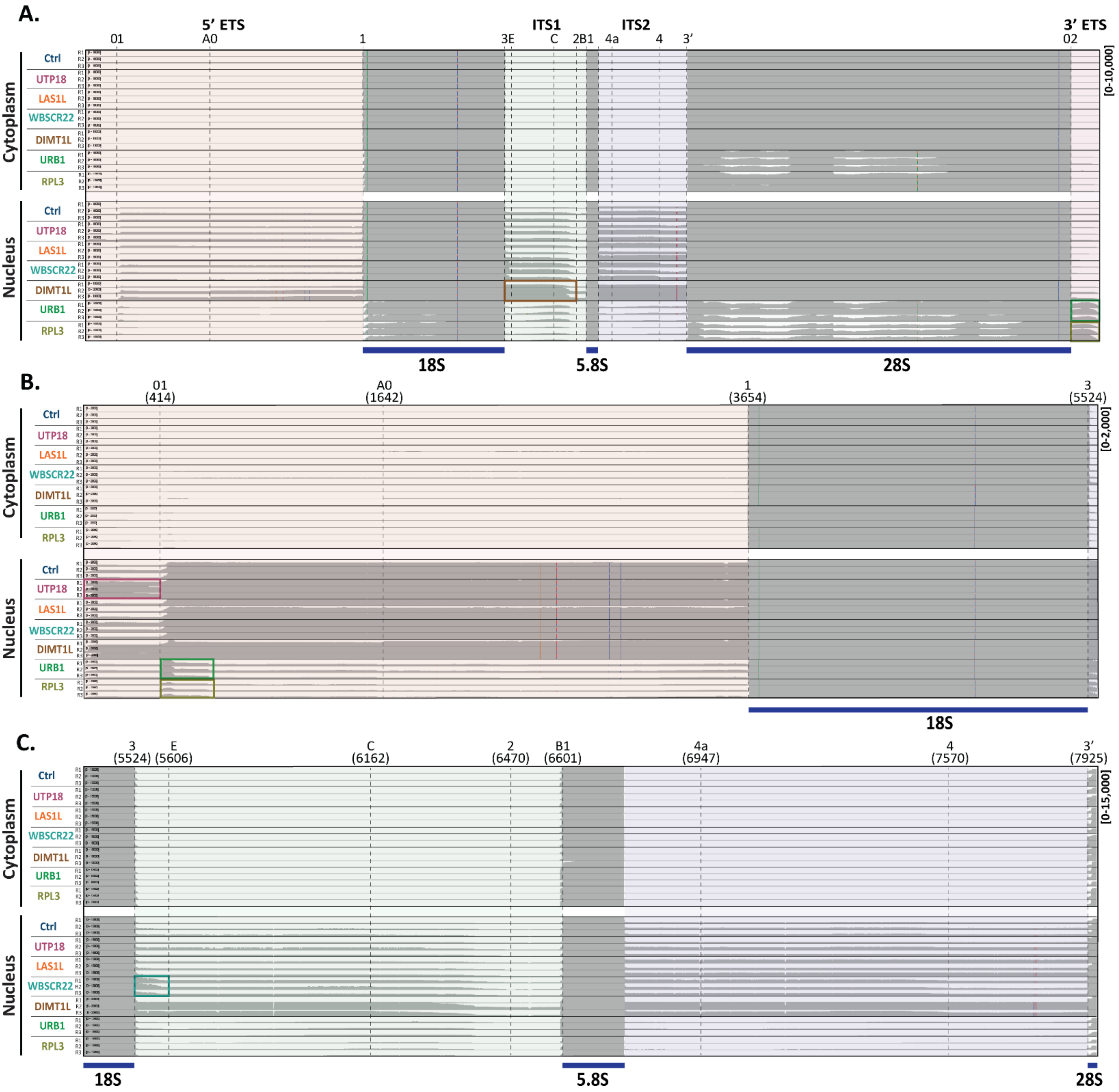


***Supplementary Figure S7. IGV snapshots of nucleus/cytoplasm in knockdown samples and Ctrl.***

**A-C**, Coverage profiles of control and knockdown conditions for UTP18, WBSCR22, DIMT1L, LAS1L, URB1 and RPL3 in nucleus/cytoplasm (n=3). The profiles are shown across the entire 47S (A), 5' ETS and 18S (B), and ITS1, 5.8S, and ITS2 (C). Data ranges were normalized across all samples to visualize the coverage profiles within the selected regions. Altered regions in the selected knockdowns are highlighted in purple (UTP18 KD), turquoise (WBSCR22 KD), brown (DIMT1L KD) and green (URB1 KD), and dark yellow (RPL3 KD).

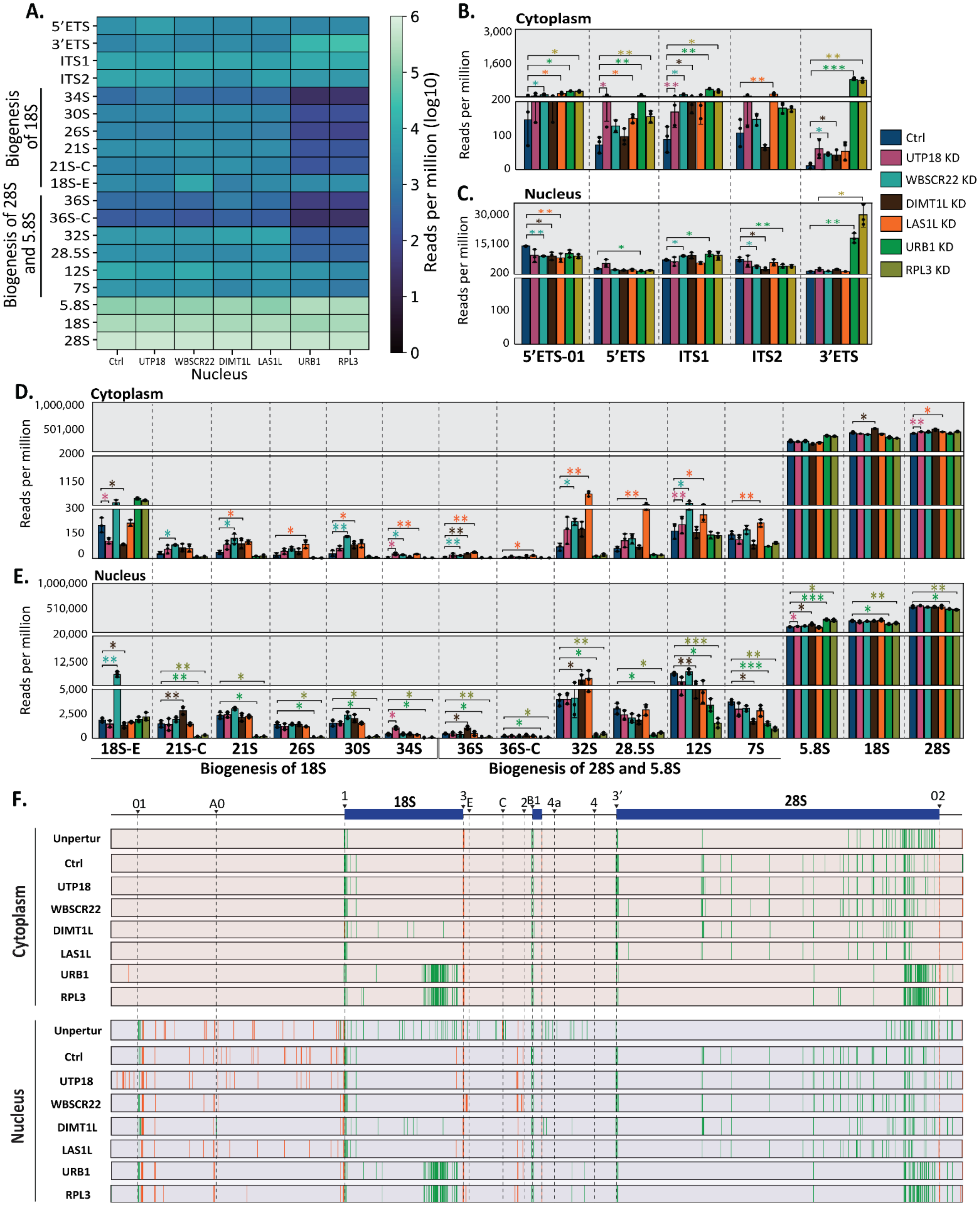


***Supplementary Figure S8. Pre-rRNA intermediate quantification and visualization of cleavages sites following processing perturbations.***

**A-B**, Quantification of pre-rRNA intermediates and mature rRNAs in cytoplasm (A) and nucleus (B) of control and knockdown samples of UTP18, DIMT1L, WBSCR22, LAS1L, URB1, and RPL3 (n=3 each).

**C-D**, Quantification of external and internal transcribed spacers in cytoplasm (C) and nucleus (D) across control and knockdown samples of UTP18, DIMT1L, WBSCR22, LAS1L, URB1, and RPL3 (n=3 each). Scores represent reads per million, while the bar plots show the means ± SD. Statistical significance was determined using a t-test between control and KD condition, with $*p < 0.05$, $**p < 0.01$, $***p < 0.001$.
**E**, Heatmap illustrating pre-/rRNA abundance in nuclear fraction of control and UTP18, DIMT1L, WBSCR22, LAS1L, URB1 knockdowns samples, represented as log10 reads per million.
**F**, Visual representation of prevalent start (green) and end (orange) cleavage sites across the 47S rRNA in cytoplasmic (top) and nuclear (bottom) samples, comparing unperturbed and control samples compared to knockdown samples of UTP18, DIMT1L, WBSCR22, LAS1L, URB1, and RPL3. Each cleavage site displayed was detected in at least two out of three replicates.

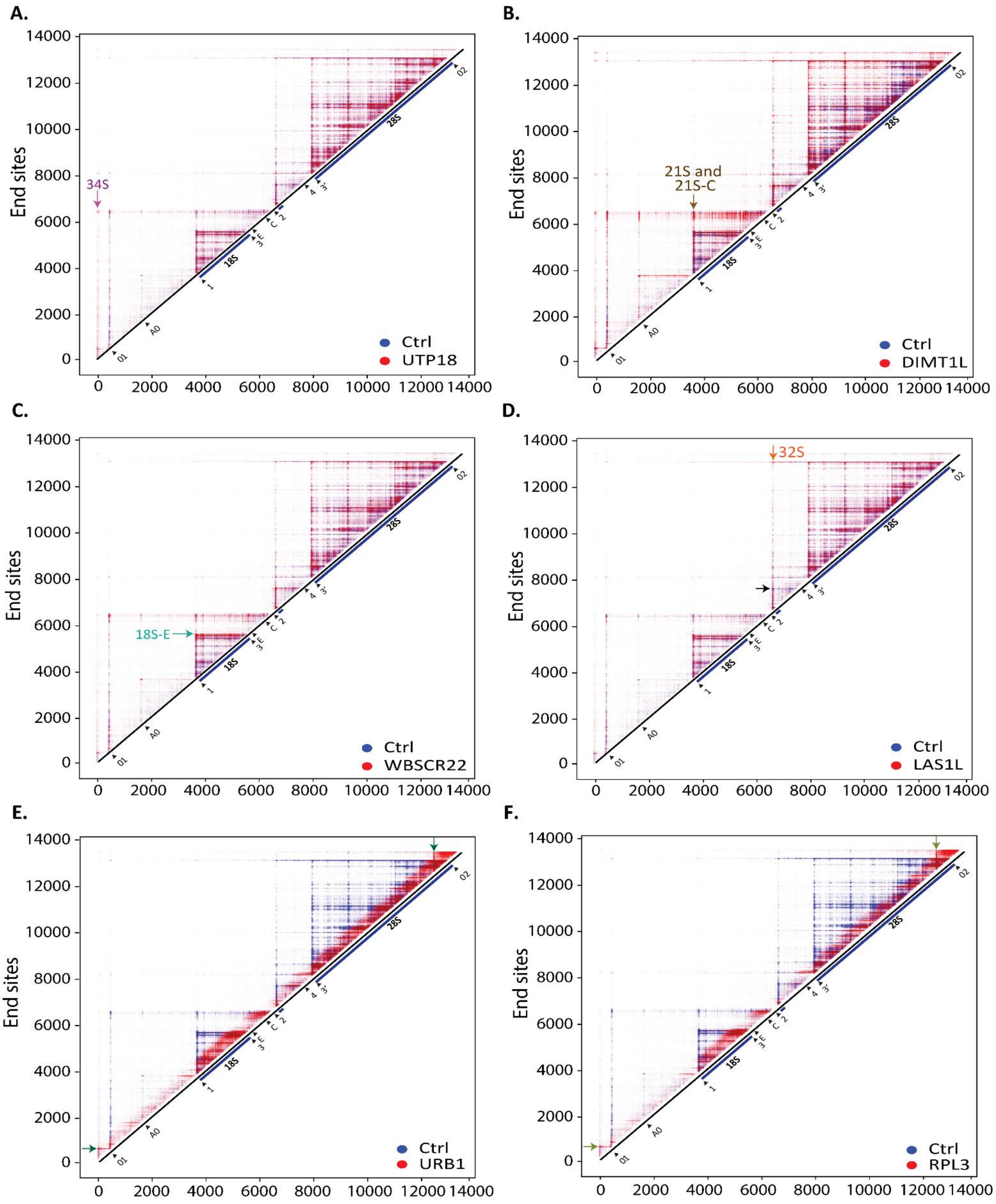


***Supplementary Figure S9. Pairwise overlay of intensity matrices of processing perturbation samples.***

**A-F**,Overalyed matrices are shown in nuclear fraction nucleus between control (blue) and designated KD conditions (red) (**A** - UTP18; **B** - DIMT1L; **C** - WBSCR22; **D** - LAS1L; **E** - URB1; **F** - RPL3). Selected intensity “hubs” associated with the effected precursors are indicated with arrows colored according to KD condition shown in Figure 4B. The black arrow in panel C marks processing site 4, the endonucleolytic site of LAS1L.

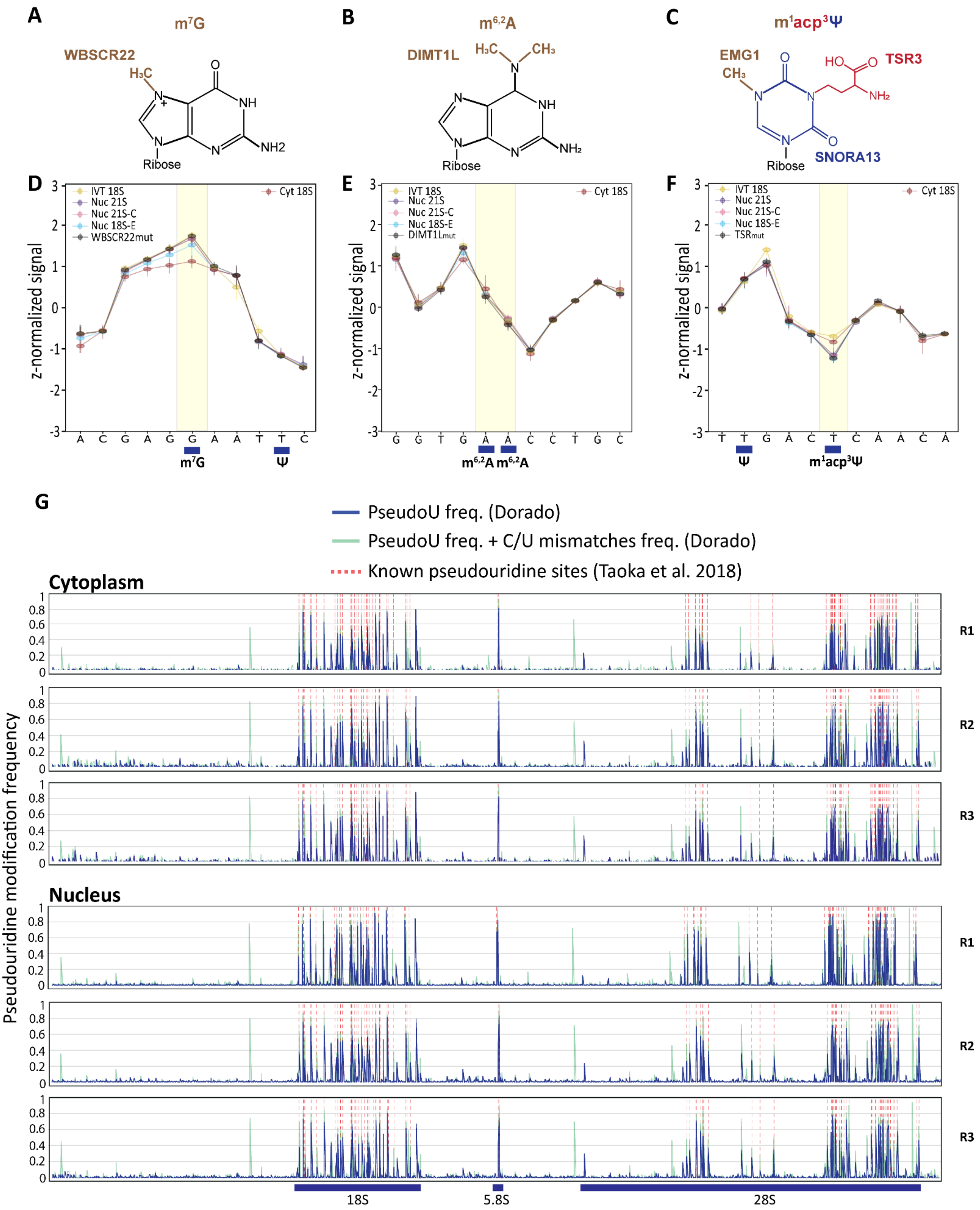


***Supplementary Figure S10. Spatio-temporal precursor-specific RNA modification and Pseudouridylation modification detection along 47S.***

**A-C,** Chemical structure of RNA modifications m7G (A), $m_2^6A$ (B) and m1acp3Ψ (C). WBSCR22[35] is a methyltransferase responsible for adding a methyl group at G[1639] on 18S rRNA, DIMT1L[35] deposits

$m_2^6A$ at $A^{1735}$ and $A^{1736}$, and TSR3[47] catalyzes the addition of α-amino-α-carboxyl-propyl (acp) following the pseudouridylation guided by SNORA13, m1 methylation via EMG1 at $U^{1248}$ on 18S rRNA.

**D-F**, The mean and standard deviation of resquiggled, z-normalized raw current signals across 11 nucleotides flanking the modification sites on 18S rRNA are shown for $m^7G$ (D), $m_2^6A$ (E), and $m^1acp^3\Psi$ (F). Data is presented for nuclear precursors (21S, 21S-C, 18S-E) and mature cytoplasmic 18S rRNA. Negative controls include IVT 18S and the corresponding mutant or knockout lines (WBSCR22mut, DIMT1Lmut, TSR3mut).

**G,** PseudoU modification frequencies in cytoplasm (top) and nucleus (bottom) across 47S rRNA landscape (n=3). Only pseudoU frequencies (Dorardo) are shown in dark blue, summed pseudoU and C/U mismatches are shown in light green and known pseudoU sites (Taoka et al., 2018) are marked by dotted red lines. Note that sites showing only C/U mismatches without corresponding pseudouridine detection are likely onlu C/U mismatches rather than pseudouridines. For further detail in Source Data 3.

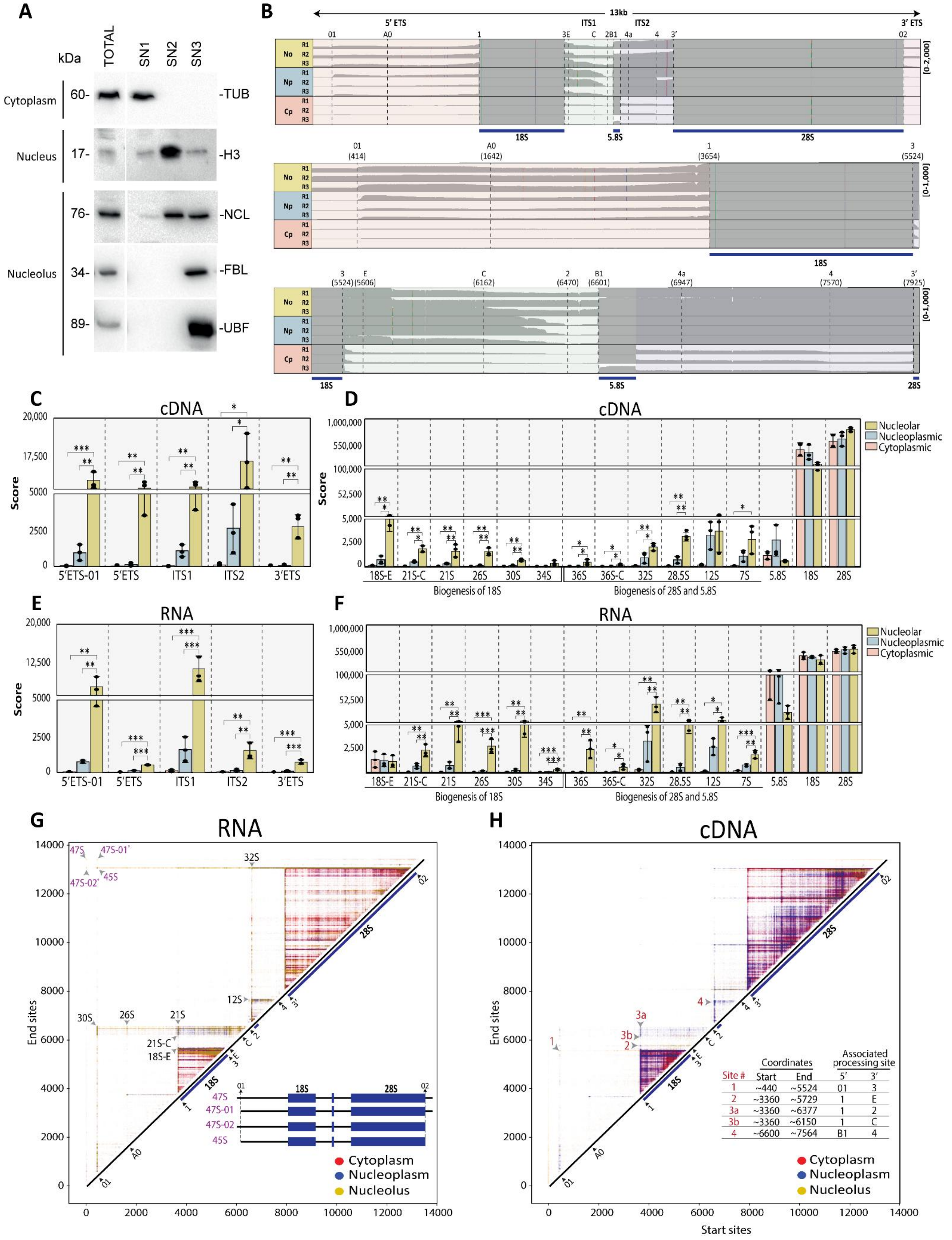

A
kDa
TOTAL
SN1
SN2
SN3
Cytoplasm
Nucleus
Nucleolus
60-
17-
76-
34-
89-
-TUB
-H3
-NCL
-FBL
-UBF
B
13kb
5' ETS
ITS1
ITS2
3' ETS
No
Np
Cp
R1
R2
R3
[0-2,000]
[0-1,000]
18S
5.8S
28S
01 (414)
A0 (1642)
1 (3654)
3 (5524)
E (5606)
C (6162)
2 (6470)
B1 (6601)
4a (6947)
4 (7570)
3' (7925)
C
cDNA
Score
5'ETS-01
5'ETS
ITS1
ITS2
3'ETS
D
cDNA
Nucleolar
Nucleoplasmic
Cytoplasmic
18S-E
21S-C
21S
26S
30S
34S
36S
36S-C
32S
28.5S
12S
7S
5.8S
18S
28S
Biogenesis of 18S
Biogenesis of 28S and 5.8S
E
RNA
F
RNA
G
RNA
47S
47S-01
47S-02
45S
32S
12S
30S
26S
21S
21S-C
18S-E
End sites
Cytoplasm
Nucleoplasm
Nucleolus
H
cDNA
Coordinates
Associated processing site
Site #
Start
End
5'
3'
1 ~440 ~5524 01 3
2 ~3360 ~5729 1 E
3a ~3360 ~6377 1 2
3b ~3360 ~6150 1 C
4 ~6600 ~7564 B1 4
Start sites

***Supplementary Figure S11. Anlaysis of pre-rRNA intermediates in cytoplasm, nucleoplasm and nucleolus.***

**A**, Western blot analysis on SN1 (Cytoplasm), SN2 (Np), and SN3 (No) fractions confirming successful fractionation. β-Tubulin (TUB) was used as a cytoplasmic marker, Histone H3 (H3) as a nuclear marker, and Nucleolin (NCL), Fibrillarin (FBL), and Upstream Binding Factor (UBF) as nucleolar markers. Marker-specific enrichment in the corresponding fractions confirms the purity and integrity of the subcellular preparations.

**B**, Coverage profiles of nucleolar (No), nucleoplasmic (Np) and cytoplasmic (Cp) conditions (n=3) in cDNA data. The profiles are shown across the entire 47S template (top), 5' ETS and 18S (middle), and ITS1, 5.8S, and ITS2 (bottom). Data ranges were normalized across all samples to visualize the coverage profiles within the selected regions.

**C-F**, Quantitative comparison of cDNA (C-D) and RNA (E-F) datasets of the ETS and ITS regions (C,E) and pre-rRNA intermediates (D,F) across the nucleolar (No), nucleoplasmic (Np) and cytoplasmic (Cp) conditions (n=3) in cDNA (top) and DRS (bottom) data. One-way ANOVA followed by Tukey test for multiple comparison post hoc test, $^{*}p < 0.05$, $^{**}p < 0.01$, $^{***}p < 0.001$. Score (read per million) represents the means ± SD.

**G-H**, Comparison of overlayed intensity matrices of nucleolar (No), nucleoplasmic (Np) and cytoplasmic (Cp) conditions in RNA (G) and cDNA (H). Key intensity hubs and their associated precursors are marked with arrows. In G, the top left (purple) highlights the positions of 47S, 45S, and the 47S variants (47S-01 and 47S-02). In H, the tables provide detail of the site numbers of selected "hubs", associated start and end coordinate (on 47S) and the respective 5' and 3' processing site.

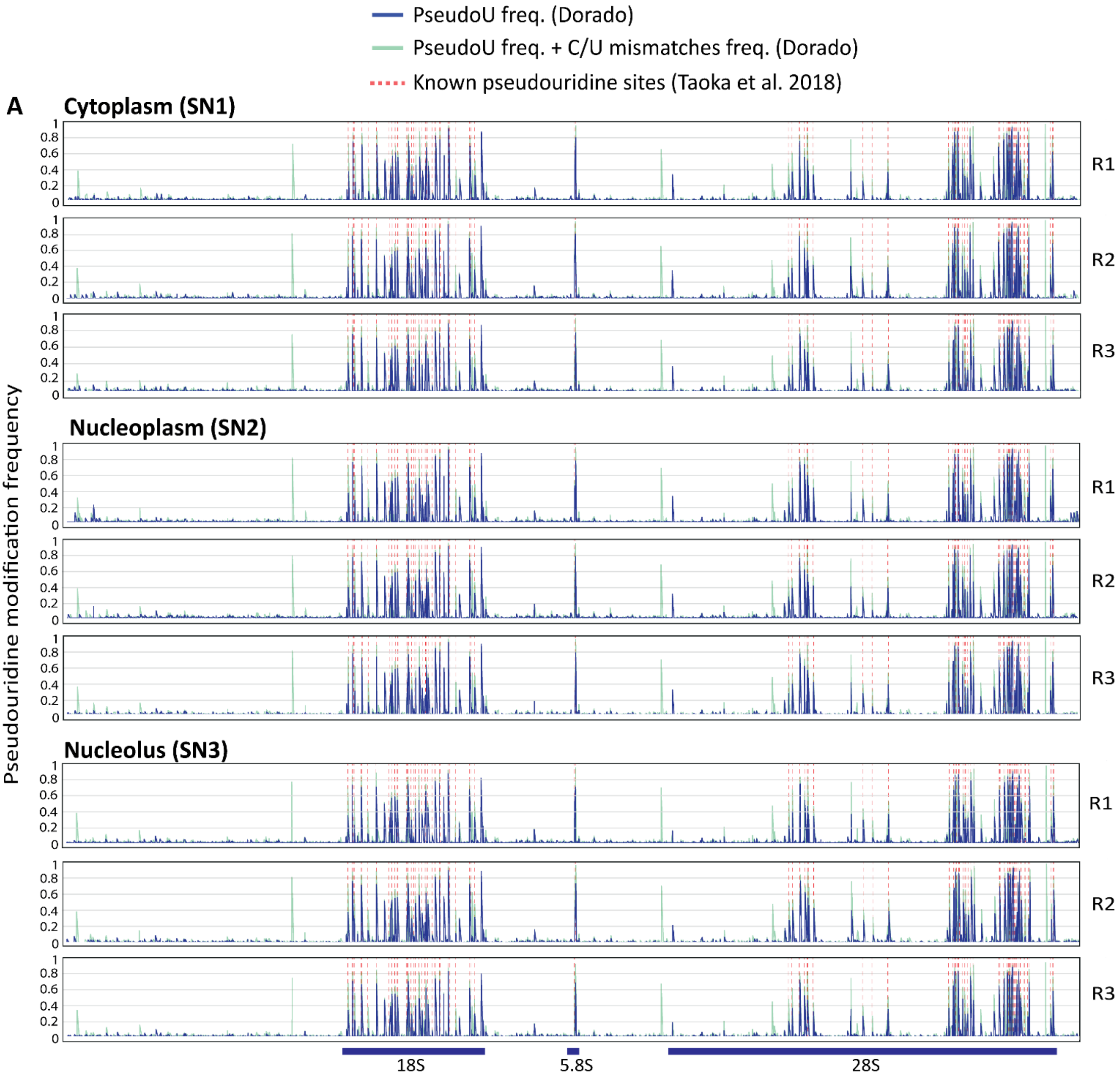


***Supplementary Figure S12. Detection of pseudouridylation across full-length 47S pre-rRNA in cytoplasmic, nucleoplasmic, and nucleolar fractions using NanoRibolyzer.***

**A,** PseudoU modification frequency in cytoplasmic (top), nucleoplasmic (middle) and nucleolar (bottom) fractions across 47S rRNA (n=3). Only pseudoU frequencies (Dorado) are shown in dark blue, summed pseudoU and C/U mismatches are shown in light green and known pseudoU sites (Taoka et al., 2018) are marked by dotted red lines. Note that sites showing only C/U mismatches without corresponding pseudouridine detection are likely only C/U mismatches rather than pseudouridines. For further detail in Source Data 3.

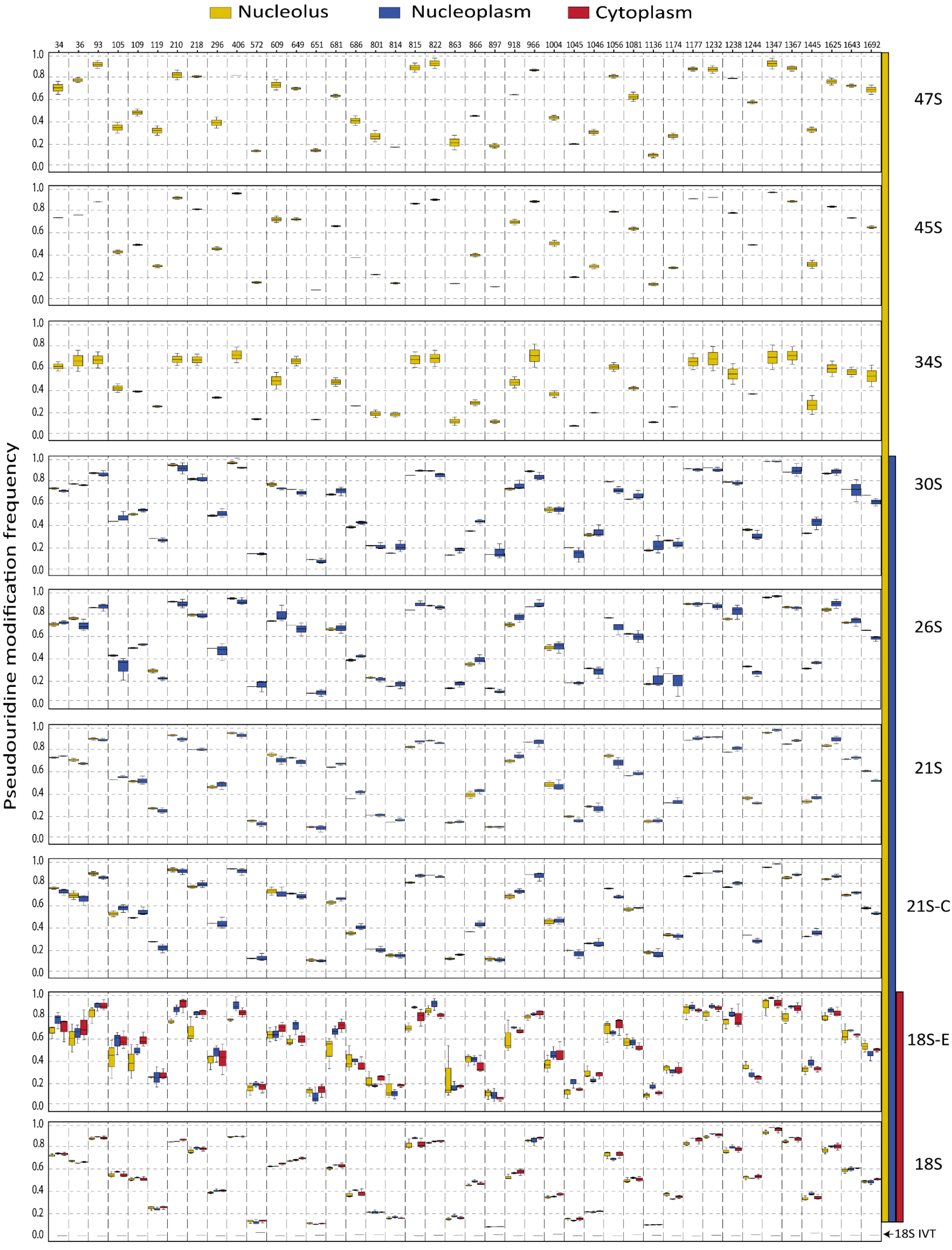

Nucleolus
Nucleoplasm
Cytoplasm
34 36 93 105 109 119 210 218 296 406 572 609 649 651 681 686 801 814 815 822 863 866 897 918 966 1004 1045 1046 1056 1081 1136 1174 1177 1232 1238 1244 1347 1367 1445 1625 1643 1692
Pseudouridine modification frequency
1.0
0.8
0.6
0.4
0.2
0.0
47S
45S
34S
30S
26S
21S
21S-C
18S-E
18S
18S IVT

***Supplementary Figure S13. Pseudouridine modification frequency across 18S rRNA during ribosome biogenesis.***

Pseudouridine modification frequencies are shown for individual precursors during 18S rRNA maturation, spanning the nucleolar (yellow), nucleoplasmic (blue), and cytoplasmic (red) fractions. Each row represents a distinct rRNA precursor, ordered from earliest (47S) to mature 18S. The x-axis denotes the pseudouridine sites along 18S rRNA (positions indicated above). The 18S IVT serves as an unmodified negative control.

**Supplementary Tables**

**Table S1.** Normalized read count per million (CPM) of Xist and Malat1 transcripts in cytoplasmic, whole cell, and nuclear fractions (n=3). Fold change (FC) between the conditions is shown above each comparison.

**Normalized counts**

| Geneid | ENSG00000229807.13 | ENSG00000251562.10 |
|---|---|---|
| gene_name | XIST | MALAT1 |
| Cytoplasm1_cpm | 395.34 | 20.81 |
| Cytoplasm2_cpm | 302.24 | 65.70 |
| Cytoplasm3_cpm | 408.33 | 72.06 |
| Cell1_cpm | 4080.08 | 464.70 |
| Cell2_cpm | 2407.99 | 210.92 |
| Cell3_cpm | 2060.61 | 281.88 |
| Nucleus1_cpm | 69789.40 | 7540.67 |
| Nucleus2_cpm | 64370.22 | 6857.48 |
| Nucleus3_cpm | 49221.52 | 7795.12 |

**Foldchange(FC)**

| gene | XIST | MALAT1 |
|---|---|---|
| nuc_vs_cell | 21.45141 | 23.17845 |
| nuc_vs_cyt | 165.8204 | 139.9601 |
| cell_vs_cyt | 7.730049 | 6.038373 |

### Supplementary Notes

#### Supplementary Note S1. *PseudoU basecaller*

Our data reliably reproduced and identified all reported pseudouridines (pseudoU) sites from the reference study by Taoka et al. (2018). However, accurate quantification of pseudoU levels remains a challenge. Multiple studies have noted that pseudoU modifications can frequently produce U-to-C mismatches during sequencing (Begik et al., 2021; Tavakoli et al., 2023; Makhamreh et al., 2024; Hewel et al., 2024, Schartel et al. 2024). This issue complicates pseudoU quantification, as it relies on a two-step analysis: first, basecalling from pod5 files identifies nucleotide composition (i.e. U), and then Remora assesses modification status to confirm pseudoU sites within basecalled U. Yet, if some pseudoU are misbasecalled as C, they bypass the Remora model, resulting in underrepresentation of pseudoU. A potential approach to more accurately estimate pseudoU levels is to utilize the U-to-C mismatch ratio as an indicator of potential pseudoU. By summing up the U-to-C mismatch rate with the identified pseudoU, we can derive a more realistic, though not precise, estimate of pseudoU presence at a given site (Hewel et al., 2024). With continued improvements in basecalling models, these limitations are likely to be resolved, enabling more reliable and easily generated datasets for studying RNA modifications.

#### Note S2. *Differences between DRS and cDNA-sequencing*

In this work, we employ both cDNA sequencing and direct RNA sequencing (DRS) to capture pre-rRNAs and mature rRNAs. cDNA sequencing enables cost-effective multiplexing of samples, allowing for the inclusion of replicates for statistical analysis. In contrast, DRS was primarily utilized for detection RNA modifications at the signal level and using pseudoU basecaller. Additionally, we employed a recently developed barcoding strategy for DRS (SeqTagger), allowing us to multiplex up to four samples per flow cell, a capability that was not available until recently (Pryszcz et al., 2025). Thus, this approach also provides a cost-effective solution for DRS, with multiple replicates within the same flow cell. While cDNA and DRS approaches yielded largely reproducible results (**Figure 5H**), slight variations were observed, likely due to differences in library preparation of each method. For instance, for cDNA libraries, Maxima H Minus RT (Thermo Fisher) was used at 42°C (ONT protocol) to synthesize first-strand cDNA with a 5′ CCC overhang, enabling barcoding and high-throughput sequencing. In contrast, DRS employed the recently developed Induro RT (NEB), a thermostable enzyme capable of reverse transcription at 55°C, improving the yield, read length, and resolution of structured RNAs like pre-rRNA (Zeglinski et al., 2024). This enhanced performance likely enabled better capture of long precursors such as 47S (~13 kb), 45S, and novel variants 47S-01 and 47S-02, particularly evident in unsupervised approach. Although Induro RT can generate very long cDNA fragments, it has not yet been coupled with second-strand synthesis for barcoded, high-throughput cDNA sequencing. Future work should explore this potential to further expand sequencing capabilities.

#### References


1. Begik, O., Diensthuber, G., Liu, H., Delgado-Tejedor, A., Kontur, C., Niazi, A. M., Valen, E., Giraldez, A. J., Beaudoin, J. D., Mattick, J. S., & Novoa, E. M. (2022). Nano3P-seq: transcriptome-wide analysis of gene expression and tail dynamics using end-capture nanopore cDNA sequencing. *Nature Methods 2022 20:1*, *20*(1), 75–85.

2. Tavakoli, S., Nabizadeh, M., Makhamreh, A., Gamper, H., McCormick, C. A., Rezapour, N. K., Hou, Y. M., Wanunu, M., & Rouhanifard, S. H. (2023). Semi-quantitative detection of pseudouridine modifications and type I/II hypermodifications in human mRNAs using direct long-read sequencing. *Nature Communications*, *14*(1), 334.

3. Makhamreh, A., Tavakoli, S., Fallahi, A., Kang, X., Gamper, H., Nabizadehmashhadtoroghi, M., Jain, M., Hou, Y. M., Rouhanifard, S. H., & Wanunu, M. (2024). Nanopore signal deviations from pseudouridine modifications in RNA are sequence-specific: quantification requires dedicated synthetic controls. *Scientific Reports 2024 14:1*, *14*(1), 1–13.

4. Hewel, C., Hofmann, F., Dietrich, V., Wierczeiko, A., Friedrich, J., Jenson, K., Mündnich, S., Diederich, S., Sys, S., Schartel, L., Schweiger, S., Helm, M., Lemke, E. A., Linke, M., &

Gerber, S. (2024). Direct RNA sequencing (RNA004) allows for improved transcriptome assessment and near real-time tracking of methylation for medical applications. *BioRxiv*, 2024.07.25.605188.

5. Schartel, L., Jann, C., Wierczeiko, A., Butto, T., Mündnich, S., Marchand, V., Motorin, Y., Helm, M., Gerber, S., & Lemke, E. A. (2024). Selective RNA pseudouridinylation in situ by circular gRNAs in designer organelles. *Nature Communications 2024 15:1*, *15*(1), 1–10. https://doi.org/10.1038/s41467-024-53403-1

6. Pryszcz, L. P., Diensthuber, G., Llovera, L., Medina, R., Delgado-Tejedor, A., Cozzuto, L., Ponomarenko, J., & Novoa, E. M. (2025). Rapid and accurate demultiplexing of direct RNA nanopore sequencing data with SeqTagger. Genome Research, 35(4), 956–966. https://doi.org/10.1101/GR.279290.124/-/DC1

7. Zeglinski, K., Montellese, C., Ritchie, M. E., Alhamdoosh, M., Vonarburg, C., Bowden, R., Jordi, M., Gouil, Q., Aeschimann, F., & Hsu, A. (2024). An optimized protocol for quality control of gene therapy vectors using nanopore direct RNA sequencing. Genome Research, 34(11), 1966–1975. https://doi.org/10.1101/GR.279405.124
